# Evidence for conformational change-induced hydrolysis of β-tubulin-GTP

**DOI:** 10.1101/2020.09.08.288019

**Authors:** Mohammadjavad Paydar, Benjamin H. Kwok

**Author notes:** Corresponding author: B. H. Kwok.

## Abstract

Microtubules, protein polymers of α/β-tubulin dimers, form the structural framework for many essential cellular processes including cell shape formation, intracellular transport, and segregation of chromosomes during cell division. It is known that tubulin-GTP hydrolysis is closely associated with microtubule polymerization dynamics. However, the precise roles of GTP hydrolysis in tubulin polymerization and microtubule depolymerization, and how it is initiated are still not clearly defined. We report here that tubulin-GTP hydrolysis can be triggered by conformational change induced by the depolymerizing kinesin-13 proteins or by the stabilizing chemical agent paclitaxel. We provide biochemical evidence that conformational change precedes tubulin-GTP hydrolysis, confirming this process is mechanically driven and structurally directional. Furthermore, we quantitatively measure the average size of the presumptive stabilizing “GTP cap” at growing microtubule ends. Together, our findings provide the molecular basis for tubulin-GTP hydrolysis and its role in microtubule polymerization and depolymerization.

## INTRODUCTION

Microtubules (MTs), protein polymers of α/β-tubulin dimers, are a major component of the cell cytoskeleton. MTs provide the framework for giving a cell its shape, for intracellular transport and for segregating the chromosomes during cell division. The formation of MT polymers and the regulation of their dynamics in space and time are crucial for their functions during different phases of the cell cycle. The fundamental questions of how these polymers are formed and their dynamics controlled at the molecular level are still not fully understood.

### MT polymerization and tubulin-Guanosine triphosphate (GTP) hydrolysis

Microtubule polymerization occurs when α/β-tubulin subunits join together to form a polymer of hollow tubule [1]. Polymerization is initiated by a process called nucleation – the assembly of two or more tubulin dimers. This can happen spontaneously or build on pre-existing templates such as the γ-tubulin ring complex [2-4]. Once the nucleus is formed, more subunits can be added to form filaments and are self-organized into a hollow tubular structure. It has been well established that MT polymerization is a stochastic process, with polymers cycling between periods of growing and shrinking. This phenomenon is known as dynamic instability [5].

It was known early on that MT polymerization requires GTP [6-8] and that α/β-tubulin dimers, subunits of MTs are GTPases. In their nucleotide-binding pockets, α-tubulin contains a non-interchangeable GTP constitutively while β-tubulin has a hydrolysable and exchangeable GTP [9-11]. GTPs in β-tubulin subunits are hydrolyzed during MT polymerization and within the MT lattice in two steps: from GTP to guanosine diphosphate(GDP)-P_i_ and then to GDP [12, 13]. GTP hydrolysis is presumed to contribute to MT polymerization dynamics and regulating the catastrophe switch [14], but the underlying molecular mechanism is still poorly understood.

Early EM study showed that polymerizing ends are often blunt and straight while the depolymerizing ends are curved or tapered [15]. This led to the pleasingly simple hypothesis that GTP-tubulins are straight, and the GDP tubulins are curved. The straight conformation of GTP-tubulin facilitates the incorporation of dimers into polymer ends forming the stabilizing GTP cap and the formation of lateral bond with adjacent protofilaments. As polymerization proceeds, GTP gets hydrolyzed and the GDP-tubulins in the MT middle are constrained to the straight conformation and are inherently unstable [16, 17]. Upon removal of the stabilizing GTP cap, the GDP-tubulin dimers are exposed and the constrains are released, leading to the outward curved of the depolymerizing filament. This model is appealing because it explains most of biochemical and conformational events associated with microtubule polymerization and depolymerization. However, accumulating structural studies have found that both GTP- and GDP-tubulin dimers are curved [18-20], and that curved or tapered filaments are also observed in polymerizing microtubule ends [21-23]. In light of these findings, it is conceivable that curved GTP-tubulins are added to the polymerizing ends. As polymerizing ends grow and filaments straighten through interaction with adjacent filaments, GTP hydrolysis ensues. Recent high resolution cryo-electron microscopy (cryo-EM) studies on MT polymers stabilized with taxol or hydrolysis-resistant GMPCPP have suggested that GTP-bound tubulin stabilized the polymers by strengthening the lateral and longitudinal interactions between neighboring tubulin subunits, and that hydrolysis leads to the destabilization GDP-tubulin in the lattice [16, 17, 24]. Upon depolymerization, the previously constrained dimers return to their naturally curved state, and the energy stored in the constrained tubulin dimer is released to do mechanical work. While this revised model is logical and sound, it is still puzzling as to what triggers GTP hydrolysis and why since GTP hydrolysis per se is not needed for the addition of GTP-tubulin subunit at MT ends.

### GTP cap and MT stability

Being enriched in GTP-bound tubulin dimers, the growing end of microtubules is often referred to as “GTP cap”; this structure is thought to stabilize MTs [12, 25]. It has also been proposed that some GTP-bound tubulin dimers may exist in MT middle (GTP islands), due to incomplete GTP hydrolysis during polymerization [12, 25]. Evidence for this proposal has been obtained indirectly using a conformation-specific antibody [26]. The islands of GTP-bound tubulin have been speculated to be responsible for MT rescue, switching MTs from shortening to growing [26, 27]. The length/size of the GTP cap has been a mystery and controversial for decades. Earlier studies suggested the GTP cap size could be as small as a single GTP-tubulin layer [1, 28, 29]. This is, however, inconsistent with recent findings by indirect probing with MT end-binding (EB) proteins, which have been shown to bind preferentially to GTP-bound tubulin [30-33]. There have been efforts using this feature to determine the GTP cap size *in vivo*, and some have reported it to contain hundreds of tubulin dimers (~750 dimers spread over ~55 rows) [34, 35]. It has been proposed that the length of EB comet tail is closely linked to the growth rate and stability of MTs [31, 36, 37], strengthening the idea that the GTP cap stabilizes the MT polymers. Curiously, in a more recent study, the length of outwardly curved MT ends was measured to be about ~40-80 nm *in vitro* and also in cells from different species, using EM tomography [22]. It is not known, however, whether the curvature is linked to the nucleotide-bound state of tubulin dimers associated with the protofilament ends.

### MT dynamics in cells

In cells, MT dynamics are largely regulated by MT associated proteins (MAPs). Tubulin post-translational modifications have also been shown to regulate different MT features, including their dynamics, by altering their binding affinity to MAPs [38, 39]. Some of these proteins promote MT polymerization/stabilization, like Stu2/XMAP215/Dis1 family polymerases, which contain multiple TOG domains that bind αβ-tubulin dimers [40]. Another member of polymerizing MAPs is TPX2, which is known as an MT catastrophe suppresser [41]. On the other hand, MT depolymerizing MAPs are also important in controlling MT dynamics. Members of Kinesin-8 family are MT destabilizing motors that move towards the plus-end of MTs. The depolymerization mechanism of Kinesin-8 motor proteins, however, varies between different species [42-44]. In contrast, kinesin-13 family members (KIF2A, KIF2B and KIF2C/MCAK) are major catastrophe factors that actively remove subunits from MT ends [45, 46]. These proteins hydrolyze adenosine triphosphate (ATP) when they encounter tubulin dimers, free or in the polymers. However, their depolymerizing function only happens at the ends of MTs, where they use the energy derived from ATP hydrolysis to dissociate tubulin dimers [47, 48]. Our structural study shows each kinesin-13 monomer can bind to two tubulin dimers in tandem and bend both the intra- and inter-dimer curvature [49]. We postulate that this extreme bending of tubulin dimers by kinesin-13 ultimately leads to their dissociations from protofilament ends.

We report here that kinesin-13 proteins induce a conformational change in tubulin dimers, which in turn triggers the hydrolysis of the exchangeable GTP on β-tubulins. We provide experimental evidence to show that it is the conformational change of tubulin that leads to GTP hydrolysis and not the other way around. We further demonstrate that this structural change of tubulin dimer has to be directional, from a more curved to a straighter conformation and not vice versa, to trigger GTP hydrolysis. Furthermore, the same mechanism also occurs during tubulin polymerization. Finally, we provide a molecular account for the occurrence of GTP-tubulin associated MT ends, often termed the “GTP cap”, and quantitatively measure its average length. In sum, our work presented here reveals the inner workings of tubulin biochemistry: the molecular relationship between structural change and nucleotide hydrolysis.

## RESULTS

### Tubulin-GTP hydrolysis can be triggered by binding and unbinding of kinesin-13 proteins

The presence of GDP in β-tubulin, in our previous structural study of the KIF2A-tubulin ternary complex [49], prompted us to determine if the hydrolysis of tubulin GTP occurred spontaneously during the sample preparation and crystallization processes, or if it was induced by forming the complex with KIF2A-NM (neck+motor; amino acids 153–553). To address this directly, we first used a fluorescence-based assay to detect the presence of GDP over time when tubulin dimers either were incubated alone or with KIF2A-NM in the presence of AMP-PNP and DARPin (Designed Ankyrin Repeat Protein, a polypeptide that binds to β-tubulin and prevents MT polymerization). Our data showed that tubulin-GTP turns over very slowly on its own (with a rate of 0.0003 S^−1^), but the hydrolysis rate increased significantly when incubated with KIF2A-NM in the presence of AMP-PNP and DARPin (with an initial rate of 0.16 S^−1^,**Fig. 1A**). Note that tubulin dimers incubated with AMP-PNP-bound KIF2A alone or with DARPin alone did not promote GTP hydrolysis. From our previous study [49], we knew that DARPin could compete with the neck region of KIF2A-NM for binding to β-tubulin and thereby release KIF2a-NM from the complex, facilitating GTP turnover. Therefore, excess DARPin should disrupt the complex formation and suppress GTP hydrolysis. Indeed, DARPin titration in the same experimental setting yielded the result that is consistent with this prediction (**Fig. 1B**).

**Fig. 1.**
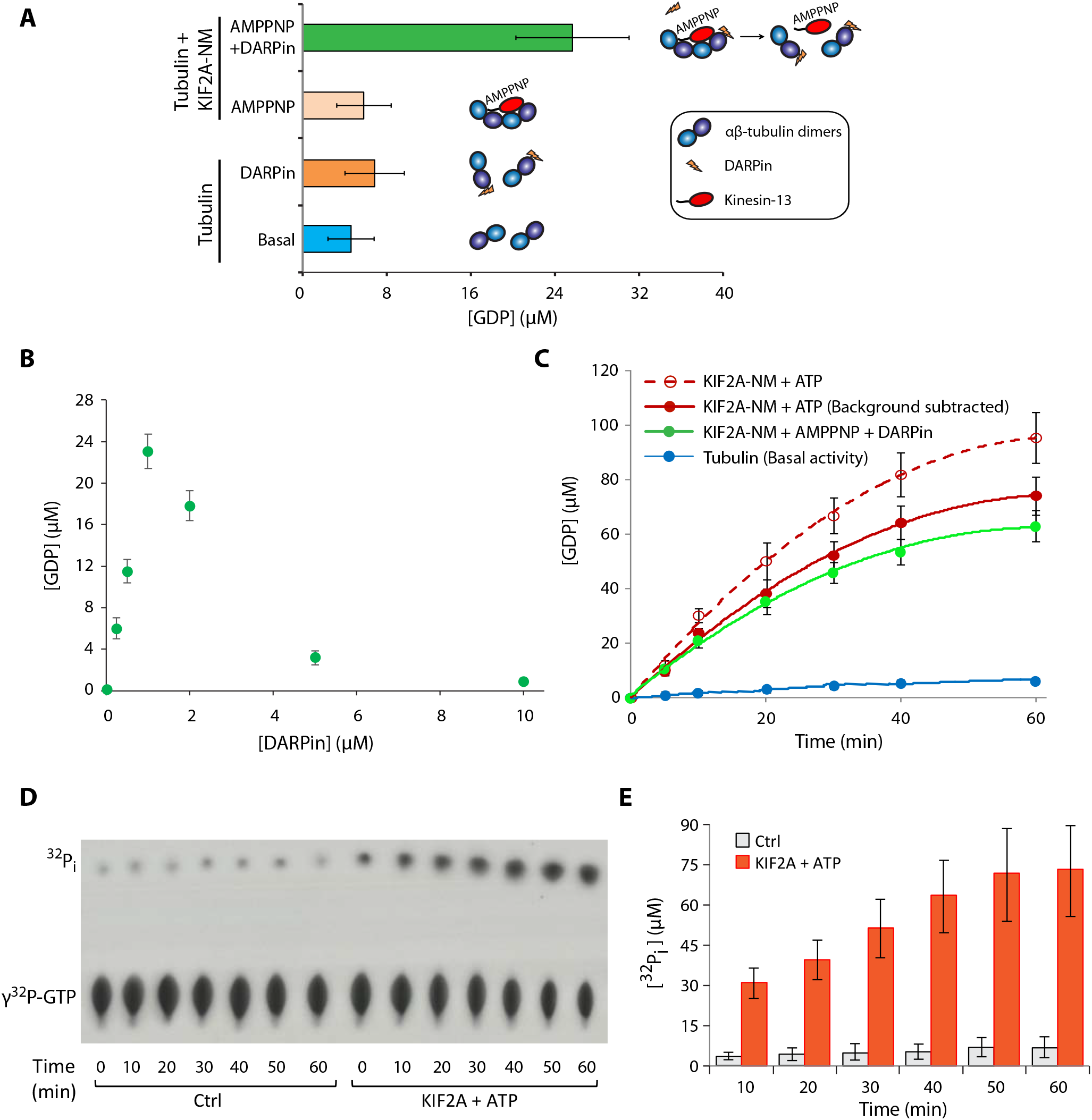
Binding dynamics of the Kinesin-13 KIF2A with tubulin dimers trigger β-tubulin-GTP hydrolysis. **(A)** Tubulin-GTP hydrolysis measured by GDP production (ProFoldin GDP detection assay) when tubulin dimers (4 μM) and GTP (0.2 mM) were incubated alone, with DARPin or in the presence of KIF2A-NM (200 nM) with AMP-PNP or with AMP-PNP and DARPin for 15 minutes at room temperature. **(B)** To determine the effect of DARPin, tubulin-GTP hydrolysis measured by GDP production when tubulin-GTP dimers were incubated in the presence of KIF2A-NM and AMP-PNP with different concentrations of DARPin, under the same condition as in (A). **(C)** Time-course experiments comparing the tubulin-GTP turnovers between tubulin dimers that were incubated in the presence of KIF2A-NM with AMP-PNP and DARPin and those with KIF2A-NM with ATP under the same condition as in (A). For (A-C), the level of GDP was measured using a GDP detection assay (ProFoldin). **(D)** A time course experiment in which basal tubulin-GTP hydrolysis was measured and compared with in KIF2A-NM-induced GTP hydrolysis (in the presence of ATP) using a γ^32^P-GTP radio-labeled GTPase assay under the same experimental setting as described as in (C). γ^32^P-GTP and ^32^P_i_ were resolved by thin layer chromatography and exposed to a film. A representative autoradiogram is shown. **(E)** Quantification of ^32^P_i_ of each time point in the time course experiment shown in (D). All the data shown represent an average of at least three independent experimental runs. Error bars represent standard deviations. (ns (not significant): p>0.05; *p≤0.05; **p≤0.01; ***p≤0.001, by Student’s *t*-test)

Our results showed that GTP hydrolysis required the associated tubulin dimers to be released from AMP-PNP-bound KIF2A-NM (i.e. bound only with DARPin, which disrupts the 1:2 KIF2A:tubulin dimer complex, as previously shown [49]). This suggests that the bending of tubulin dimers by KIF2A-NM per se does not trigger GTP hydrolysis; instead it is the return of free tubulin dimers to their native conformation that promotes the activity. If this assertion is correct, ATP hydrolysis by KIF2A, stimulated by tubulin dimers, should also trigger GTP hydrolysis. As DARPin was used only to facilitate the crystallization of KIF2A-NM tubulin complex, it should not be needed for KIF2A or kinesin-13 proteins to interact with tubulin dimers at MT ends, there they induce MT depolymerization. To ensure that what we observed was not due to an artifact of using DARPin to force the release of KIF2A-NM from tubulin dimer, and to determine if kinesin-13-induced tubulin-GTP hydrolysis also occurs in the presence of ATP, we performed the same experiment in the presence of ATP. As anticipated, this was indeed the case (**Fig. 1C**). Our result showed that GTP turnover also occurred in the presence of ATP, as robustly as or even more robust than that with AMP-PNP and DARPin. Although the GDP detection reagent that we used has a lower sensitivity to adenosine diphosphate (ADP) (see calibration curve in **Fig. S1**), the signal is measurable and therefore should be subtracted from the total signal (**Fig. 1C**, solid and dotted red lines represent before and after background subtraction, using the determined ATP turnover rate). The results obtained with ATP were comparable to those with AMP-PNP and DARPin. To obtain further evidence on this point, we performed the same experiment using radio-labeled γ^32^P-GTP as a tracer to unambiguously distinguish GTP hydrolysis from ATP hydrolysis. This experiment showed a time-dependent increase of GTP hydrolysis (**Fig. 1D-E**), consistent with the result from the GDP detection assay (**Fig. 1C)**. As a control, we monitored the basal tubulin-GTP turnover over longer periods of time, using radio-labeled γ^32^P-GTP, which showed slow basal GTP hydrolysis of 0.0002 (s^−1^) **(Fig. S2),** also consistent with the rate measured by GDP detection assay. Monitoring tubulin-GTP hydrolysis in the presence of another kinesin-13 construct, MCAK-NM, yielded results almost identical to those with KIF2A-NM **(Fig. S3)**.

### Tubulin-GTP hydrolysis occurs during MT depolymerization via catalytic depolymerization by kinesin-13s, but not by non-catalytic means

Kinesin-13 proteins, including KIF2A and MCAK, are known to depolymerize MTs [48, 50]. We wondered if the observed induction of GTP-tubulin hydrolysis occurs during MT depolymerization when kinesin-13s encounter tubulin at MT ends. To address this, we performed kinesin-13 (KIF2A-NM and MCAK-NM)-induced depolymerization assay using MT polymers that retained their GTP-bound like state. MTs can be made with slowly hydrolyzable GTP analogs, guanosine 5-3-O-(thio)triphosphate (GTPγS) or guanylyl-(alpha, beta)-methylene-diphosphonate (GMPCPP). Hydrolysis of GTPγS produces GDP, just as GTP hydrolysis does. It has been shown that kinesin-13 proteins can depolymerize these MT polymers [51]. To determine if kinesin-13 proteins can trigger GTPγS hydrolysis upon MT depolymerization, we first generated ^35^S-labeled GTPγS MTs. To reduce background radioactivity from unpolymerized tubulin and excess ^35^S-GTPγS, we separated the ^35^S-labeled GTPγS MT polymers from the rest of the reaction mixture by ultracentrifugation through a glycerol cushion. We resuspended the radio-labeled MTs in stabilizing PIPES-based buffer and used it in a kinesin-13 mediated depolymerization reaction. We resolved the GTPγS hydrolysis via thin layer chromatography (TLC) and monitored the level of MT depolymerization by sedimentation assay. From this experiment, we could observe concomitant GTPγS hydrolysis and MT depolymerization with both KIF2A-NM and MCAK-NM (**Fig. 2A-C & Fig. S4**). This result indicates that the depolymerization of these MTs by kinesin-13s can trigger GTPγS hydrolysis. Analogous experiment with GMPCPP MTs in kinesin-13-mediated depolymerization using a phosphate detection assay yielded similar result indicating that hydrolysis of GMPCPP also occurred upon MT depolymerization (**Fig. S5**). This observation is intriguing because kinesin-13-induced MT depolymerization is even strong enough to force the hydrolysis of GTPγS or GMPCPP, which normally does not occur during the polymerization process. This raises the possibility that this unusual hydrolysis occurs either when MT undergoes depolymerization, or only when conformational change is specifically induced by kinesin-13 proteins. To distinguish these two possibilities, we sought to depolymerize MTs by alternative means. Exposure to Ca^2+^ or cold temperature is known to cause MT depolymerization, including those MTs formed with GTPγS or GMPCPP [23, 52, 53]. For this, we set up depolymerization reactions using ^35^S-labeled GTPγS MTs. Interestingly, both Ca^2+^ and cold treatments did not trigger GTPγS hydrolysis (marked by the lack of ^35^S-labeled P_i_ release, **Fig. 2D top & Fig.-2E**) despite their high efficiency in depolymerizing MTs (indicated by tubulin dimer release in supernatant **Fig. 2D bottom & Fig.2F**). Together, these experiments indicate that the depolymerization of MTs induced by kinesin-13 proteins, but not by calcium or cold treatment, causes a conformational change of tubulin dimers severe enough to trigger hydrolysis of GTPγS or GMPCPP, and by inference, GTP as well.

**Fig. 2.**
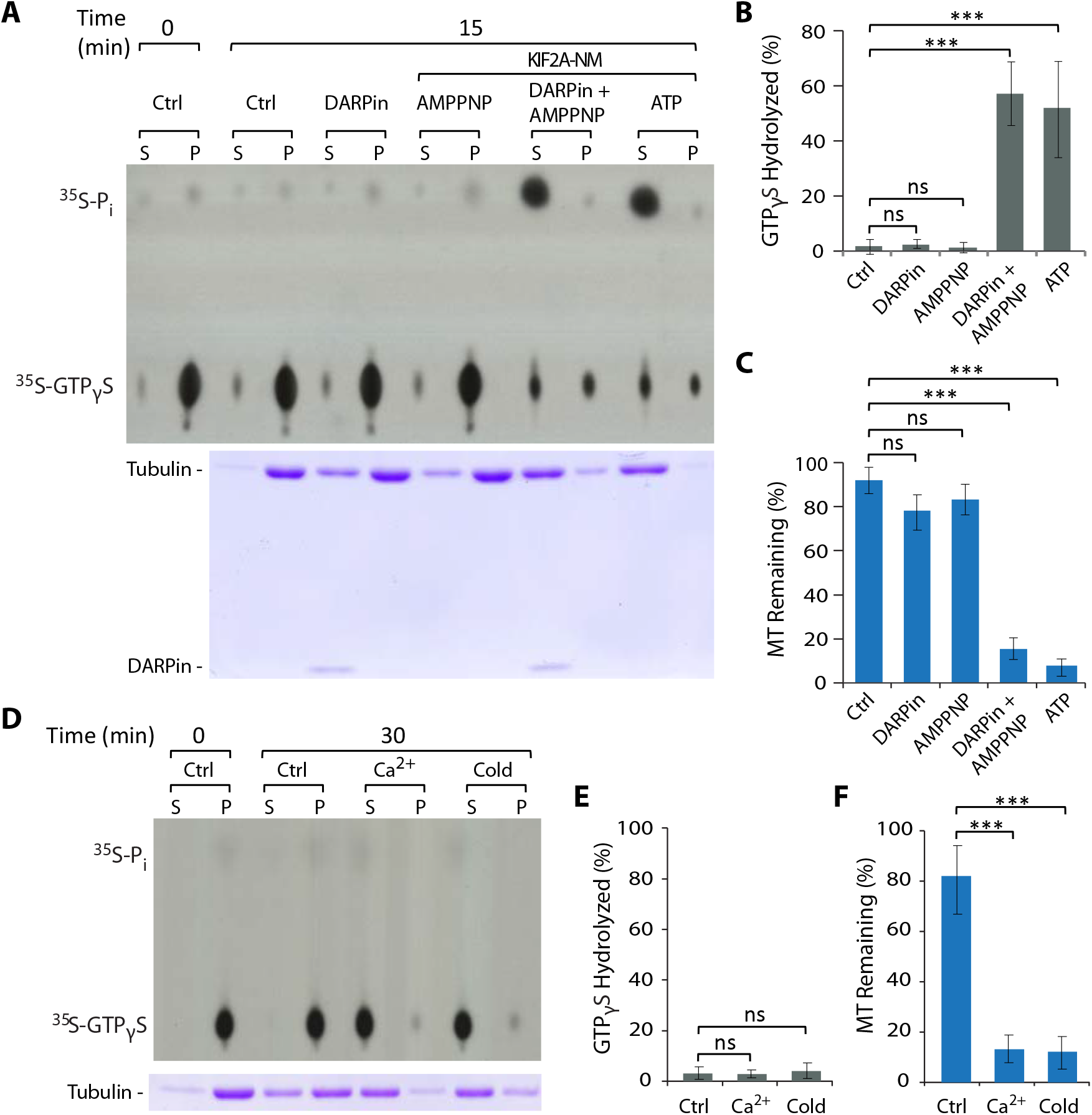
Kinesin-13 mediated MT depolymerization, unlike non-catalytically induced depolymerization, triggers tubulin-GTP hydrolysis. **(A)** MT depolymerization assay was set up using ^35^S-GTPγS-labeled MTs alone (Ctrl) or in the presence of DARPin alone, with KIF2A-NM and AMP-PNP, with KIF2A-NM, AMP-PNP and DARPin, or with KIF2A-NM and ATP. Reactions were carried out at room temperature for 15 minutes. Samples containing ^35^S-GTPγS and ^35^S-P_i_ were resolved by thin layer chromatography (TLC) and radioactivity was detected by exposure to a film. A representative autoradiogram was shown on the top panel. The level of MT polymers was monitored at the 15-minute time point using an ultracentrifugation-based sedimentation-based assay. Samples from the supernatant (S) and pellet (P) fractions were resolved by SDS-PAGE and the gel stained by Coomassie blue. A representative gel is shown on the bottom panel. MTs were used at 2 μM, KIF2A-NM at 50 nM and DARPin at 1μM. **(B-C)** Quantification of data from experiments shown in (A). B, data from autoradiograms; C, data from Coomassie blue stained gels. **(D-F)** Depolymerization of ^35^S-GTPγS-labeled MTs at room temperature (ctrl), or by treatment of cold temperature (4°C) or Ca^2+^ (1 mM) for 30 minutes. Samples were processed, quantified and analyzed in the way as those shown in (A-C). A representative autoradiogram and a Coomassie Blue stained gel were shown in (D), and their quantifications were shown in (E) and (F), respectively. All data represent the average of at least 3 independent experimental sets. Error Bars, S.D. (ns (not significant): p>0.05; *p≤0.05; **p≤0.01; ***p≤0.001, by Student’s *t*-test)

### Interdependent relationship between kinesin-13 ATPase rate and tubulin-GTP turnovers

From our previous study [49], we know that each kinesin-13-NM motor molecule can form a complex with two tubulin dimers in tandem and changes their curvatures, but we do not know whether this interaction triggers hydrolysis of both GTP molecules or just one. Our crystal structure suggests one of the two dimers is more curved than the other when they bind to KIF2A-NM with AMP-PNP. However, we do not have information as to whether the degrees of curvature are the same in the presence of ATP and whether the change(s) in curvature of either or both dimer(s) is (are) sufficient to trigger tubulin-GTP hydrolysis upon release. To address this, we first set up nucleotide hydrolysis experiments with KIF2A-NM or MCAK-NM and free tubulin dimers with radio-labeled α^32^P-ATP and γ^32^P-GTP tracers, and resolved samples by TLC (**Fig. 3A**). From these experiments, we could quantitatively measure how much ATP and GTP hydrolyzed in the same reaction based on the amount of released radio-labeled α^32^P-ADP and ^32^P_i_, respectively (**Fig. 3B**). We consistently observed that for each ATP hydrolyzed, there was approximately twice as much GTP hydrolyzed. In parallel experiments, we used the same set up and added both radio-labeled tracers in the same reaction mix, and we obtained the same result (**Fig. S6**). These data indicate that upon each encounter, for each ATP molecule hydrolyzed by KIF2A-NM, there is a corresponding hydrolysis of two GTP molecules from the two associated tubulin dimers. This also suggests that both tubulin dimers, within the ternary complex, undergo conformational changes that are severe enough to trigger tubulin-GTP hydrolysis upon their releases from the associated kinesin-13 protein.

**Fig. 3.**
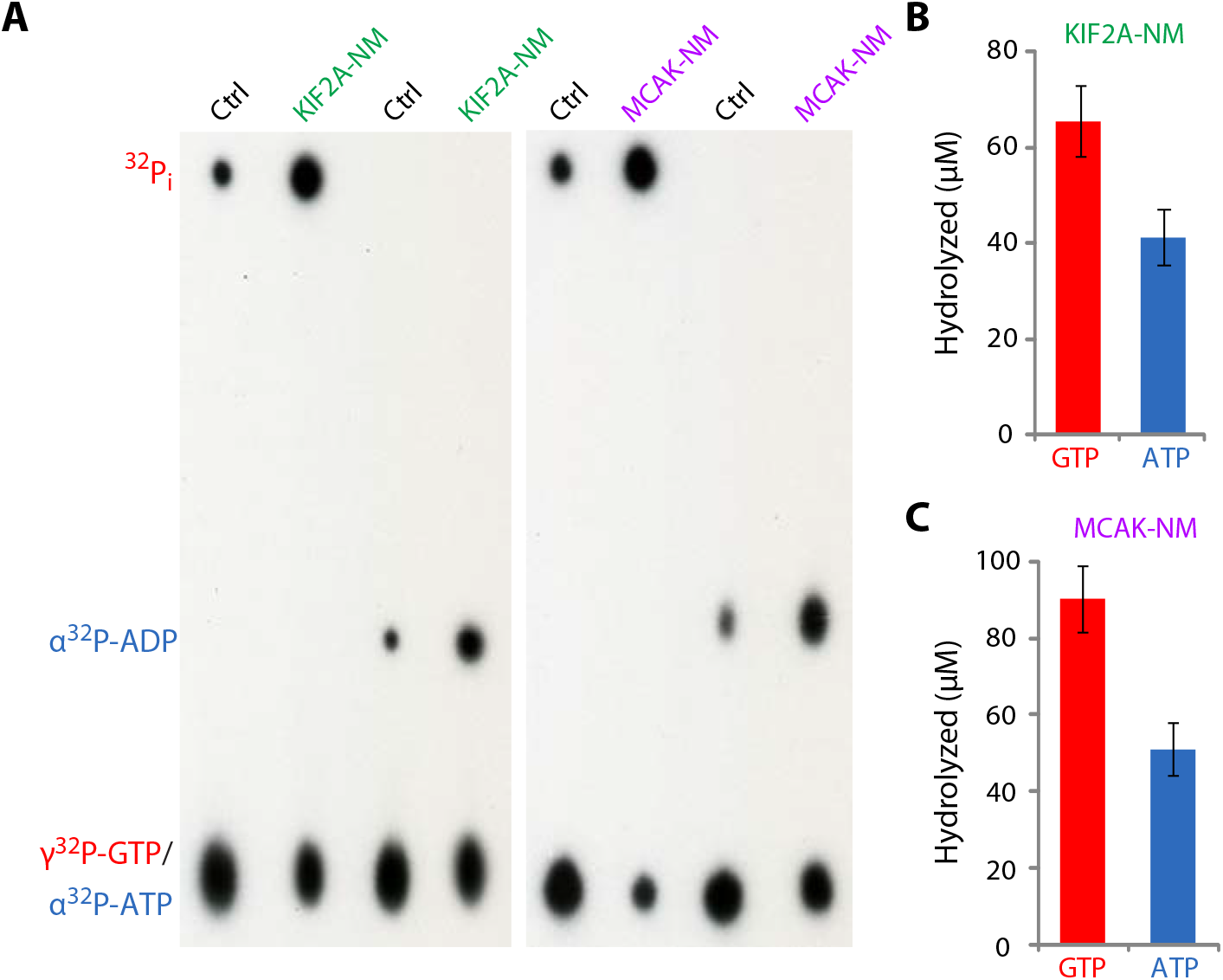
Stoichiometric relationship between kinesin-13 ATPase activity and the induced β-tubulin-GTP turnover. **(A)** Kinesin-13-induced ATP hydrolysis and tubulin-GTP turnover were monitored using radiolabeled α^32^P-ATP and γ^32^P-GTP as tracers in an enzymatic assay as described in Fig. 2. A representative autoradiogram of a TLC plate (out of 3 independent experimental runs) is shown. Tubulin dimers were used at 4 μM, KIF2A-NM and MCAK-NM at 500 nM, ATP and GTP at 200 μM. Reactions were carried out at room temperature for 10 minutes. **(B-C)** Quantification data for the experiments shown in (A). The levels of ATP and GTP hydrolysis were quantified based on the percentage of α^32^P-ADP (from α^32^P-ATP) and ^32^P_i_ (from γ^32^P-GTP) of the total amount of α^32^P-ATP and γ^32^P-GTP used in the corresponding reaction. Data represent averages of at least 3 independent experimental sets. Error Bars, S.D.

It has also been shown that MT polymers stimulate kinesin-13 ATPase activity more than free tubulin dimers [48]. We hypothesized that straighter tubulin conformations stimulate kinesin-13 ATPase activity more than the curved ones. To test this, we set up kinesin-13 ATPase assays in the presence of tubulin dimers with different guanine nucleotides (GTP, GTPγS, GDP or Apo-state). Consistent with this hypothesis, we observed that GTP- or GTPγS-tubulin dimers stimulated kinesin-13 ATPase turnovers significantly higher than those with GDP or the Apo-state using three independent detection assays (radioactivity, malachite green-based phosphate detection assay and ADP-Glo™ assay). These data are summarized in **Table 1** and **Fig. S7A-B**. This interpretation is based on the published structural data indicating that GTP-tubulin dimers (PDB code: 4DRX) are slightly straighter than GDP-tubulin (PDB code: 1SAO) [54-56]. We postulated that these differences in ATPase rate might be due to the corresponding differences in the binding affinity of tubulin dimers in different nucleotide states to kinesin-13 proteins. To test this, we set up a binding assay using tubulin dimers in different nucleotide states with either His-tagged KIF2A-NM or MCAK-NM, which could be captured by Ni-charged magnetic beads. As expected, tubulin dimers associated with GTP- or GTPγS-showed significantly higher binding affinity to both KIF2A-NM or MCAK-NM, than those with GDP- or no nucleotide (Apo) **(Fig. S7C-E)**.

**Table 1.** Stoichiometric and Interdependent relationship between Kinesin-13 ATPase rates and β-tubulin-GTP turnover rates. The rates of Kinesin-13 ATP hydrolysis and tubulin-GTP turnovers were determined using two independent methods: radio-labeled α^32^P-ATP, γ^32^P-GTP and ^35^S-GTPγS in an enzymatic assay (as shown in Fig. 3 and Fig. S6); and Malachite green-based phosphate detection assay (shown in Fig. S7). The levels of ATP hydrolysis of kinesin-13s (KIF2A-NM and MCAK-NM) were quantified for different nucleotide-bound tubulins. (* To calculate the rate of GTP hydrolysis, we had to use ATPγS instead of ATP.)

| Nucleotide state |  | Hydrolysis rate (s<sup>-1</sup>) |  |  |  |
| --- | --- | --- | --- | --- | --- |
|  |  | Hot assay |  | Malachite green |  |
|  |  | ATP | GTP | ATP | GTP |
| KIF2A-NM | Tub-GTP | 0.13 ± 0.04 | 0.23 ± 0.06 | 0.16 ± 0.04 | 0.31 ± 0.09 * |
|  | Tub-GTPγS | 0.14 ± 0.04 | 0.26 ± 0.07 | 0.18 ± 0.05 | - |
|  | Tub-GDP | 0.09 ± 0.03 | - | 0.12 ± 0.04 | - |
|  | Tub-APO | 0.11 ± 0.02 | - | 0.11 ± 0.03 | - |
| MCAK-NM | Tub-GTP | 0.15 ± 0.04 | 0.29 ± 0.06 | 0.17 ± 0.05 | 0.33 ± 0.07 |
|  | Tub-GTPγS | 0.16 ± 0.04 | 0.31 ± 0.06 | 0.18 ± 0.05 | - |
|  | Tub-GDP | 0.11 ± 0.04 | - | 0.13 ± 0.03 | - |
|  | Tub-APO | 0.10 ± 0.03 | - | 0.13 ± 0.04 | - |

Based on the ATPase rates of KIF2A-NM and the corresponding tubulin-GTPase rates summarized in Table 1, we could also observe the 1:2 ratio that we had determined earlier in Fig. 3. Together, these results demonstrated the interdependent relationship between kinesin-13 ATPase activity and tubulin-GTP turnovers as well as the associated tubulin conformational states.

### Conformational change-induced tubulin-GTP hydrolysis during MT polymerization

Our finding, that conformational change in tubulin dimers can trigger GTP hydrolysis in our kinesin-13 induced experiments, prompted us to consider the possibility that conformation-induced tubulin-GTP turnover also occurs during MT polymerization. It is known that both GTP-tubulin and GDP-tubulin dimers are more curved compared to the straight conformation of tubulin subunits within the MT lattice except for those at polymer ends. We postulated that the reason GTPγS and GMP-CPP tubulins promote MT polymerization is because they are already in a straighter conformation and that the change in conformation is not big enough to trigger hydrolysis of these slowly hydrolyzable GTP analogs. Likewise, paclitaxel promotes tubulin polymerization in the presence of GTP by straightening the associated free tubulin dimers. If this assessment is true, we should be able to observe paclitaxel-induced tubulin-GTP hydrolysis prior to polymerization. To test this possibility directly, we set up experimental conditions where tubulin-GTP and paclitaxel were both present at low concentrations to disfavor spontaneous polymerization, at 5μM and 1-4 μM respectively. To further ensure that polymerization did not occur readily, we ran the experiment at 4°C, using the GDP detection assay. As anticipated, we observed almost instantaneous conversion of GTP to GDP in the presence of paclitaxel but not at its absence (**Fig. 4A**). We also noted that the plateau of GDP accumulation was reached quickly and never exceeded the concentrations of paclitaxel used. We verified that there was no detectable level of MT polymerization under these conditions by ultracentrifugation-based sedimentation assay (**Fig. S8**). Together, these data suggest that paclitaxel binding triggers a conformational change of free tubulin dimers from their native curvature to a straighter state in the absence of polymer formation. We postulated that if this assertion is correct, paclitaxel-stabilized tubulin-GDP might also be able to form polymer. This was observed when we attempted tubulin-GDP polymerization in the presence of paclitaxel (**Fig. 4B**). The polymers that formed from tubulin-GDP are essentially the same as those formed form tubulin-GTP as assessed by their ability to sediment upon ultracentrifugation (**Fig. 4B-C**), to stimulate kinesin-13 ATPase activity (**Fig. 4E**), and by their filamentous structure observed by negative stained EM (**Fig. 4D**). Together, these results showed the following: First, tubulin-GTP hydrolysis can be resulted from paclitaxel binding and its effect on tubulin conformation. Second, tubulin-GTP hydrolysis is not an absolute requirement for MT polymerization. Third, paclitaxel-induced conformational change of tubulin dimers, likely straightening as supported by previously published data [57], can facilitate MT polymerization in the absence of tubulin-GTP (and therefore its hydrolysis as well).

**Fig. 4.**
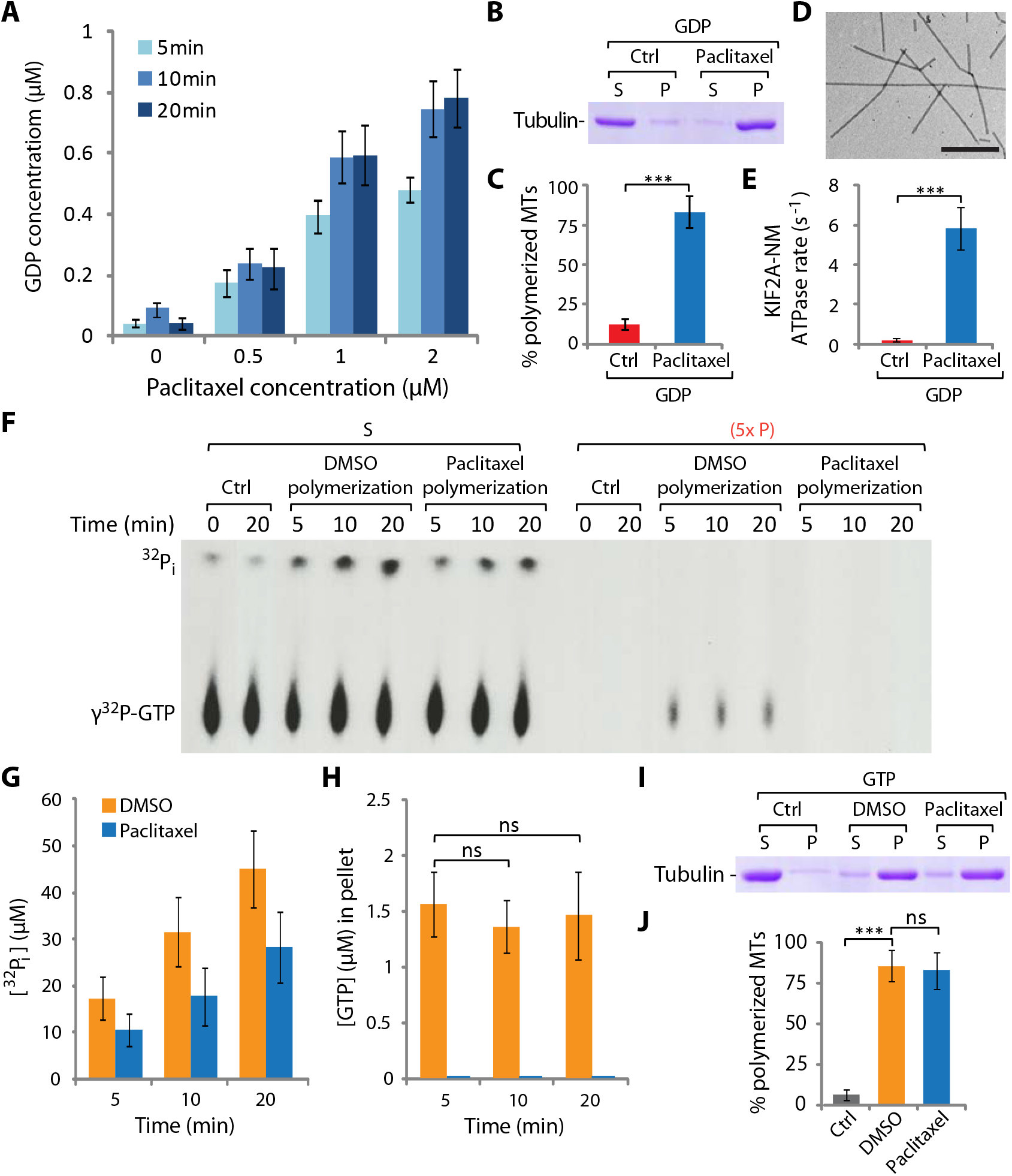
Conformational change of tubulin dimers precedes GTP hydrolysis as they incorporate into MT polymers. **(A)** The amount of tubulin-GTP hydrolyzed induced by different concentrations of paclitaxel (from 0 – 2 μM, as indicated) was measured over time (at 5, 10, 20 min) at 4°C, using a GDP detection assay. Tubulin dimers were used at 2 μM with 50 μM GTP. **(B)** MTs were polymerized using tubulin-GDP (5 μM tubulin dimers and 200 μM GDP) in the absence or presence of 20 μM paclitaxel. The polymerization reactions were carried out at 37°C for 30 min. The levels of MT polymerization were determined using an ultra-centrifugation-based sedimentation assay. Samples from the supernatant (S) and pellet (P) fractions were resolved by SDS-PAGE and the gel stained by Coomassie blue. A representative gel is shown. **(C)** The levels of polymerized MTs in (B) were quantified using ImageJ gel analysis. **(D)** Negative stained TEM images of MTs generated in (B). Scale bar, 1 μm. **(E)** The ATPase rate of KIF2A-NM in the presence of the MTs polymerized in (B) was determined using a Malachite green-based phosphate detection assay. **(F)** MTs were polymerized using 10 μM tubulin dimers in the presence of 10% DMSO or 20 μM paclitaxel (with 200 μM GTP and the presence of a radio-labeled γ^32^P-GTP tracer). GTP hydrolysis was monitored during MT polymerization by quantifying the level of γ^32^P_i_ generated, as described in Fig. 3. The unpolymerized tubulin dimers and polymerized MTs were separated into supernatant (S) and pellet (P) fractions through ultra-centrifugation. To detect the level of γ^32^P-GTP more readily, the pellet fractions were loaded at 5x equivalence of the amount of the corresponding supernatant fractions. A representative autoradiogram from three independent experimental runs is shown. **(G-H)** Quantifications of γ^32^P_i_ in the supernatant fractions (indicative of overall GTP hydrolysis) and γ^32^P-GTP in the pellet fractions (indicative of the amount of GTP on MT polymers) from the samples in F were shown in (G) and (H), respectively. **(I-J)** The levels of MT polymerization in reactions shown in (F) at 20 min were determined using a sedimentation assay. Samples were resolved by SDS-PAGE, stained and quantified in the same way as described above in (B-C). A representative Coomassie-blue stained gel (I) and the corresponding quantification (J) from three independent experiments were shown. Error bars, S.D. (ns (not significant): p>0.05; *p≤0.05; **p≤0.01; ***p≤0.001, by Student’s *t*-test)

### Evidence for tubulin-GTP at MT ends – the “GTP Cap”

Our finding that conformational change of tubulin dimers, namely its straightening, leads to GTP hydrolysis, prompted us to hypothesize the following: During MT polymerization, as tubulin dimers straighten and incorporate into protofilaments, the conformational change activates the GTPase activity of β-tubulin. In this scenario, the only tubulin dimers that remain GTP-bound are those at the polymerizing MT ends. This model predicts that during MT polymerization once nucleation has been established, the level of GTP-tubulin at MT ends should remain at similar level as the number of polymerizing ends will stay relatively constant. To test this, we carried out a time course experiment with tubulin in the presence of GTP and γ^32^P-GTP tracer, followed by ultracentrifugation-based sedimentation to separate the polymerizing MTs and unpolymerized tubulin dimers. Indeed, we observed that while the net hydrolysis of GTP increased with time as more and more polymers formed (**Fig. 4F, left**), the amount of GTP in the polymers (i.e. pellets in sedimentation assay) was constant throughout the polymerization process (comparing 5, 10, 20 min time points in **Fig. 4F, right**). For comparison, we also performed the same experiment with paclitaxel. As expected, paclitaxel eliminated GTP incorporation at MT ends (**Fig. 4J**). We also observed that the overall GTP hydrolysis in the presence of paclitaxel was lower than that of polymerization enhanced by DMSO, where growing and shrinking of MTs occurred more readily and continuously (**Fig. 4G-I**). This result is consistent with the understanding that paclitaxel favors MT polymerization, but suppresses its dynamics (and hence lower overall GTP hydrolysis, marked by lower inorganic phosphate (Pi) release than the corresponding DMSO polymerization at all time points, **Fig. 4I**).

Results from our polymerization experiments predicted that if we started with more polymerization nuclei or smaller/shorter nuclei (and therefore more nuclei at the same concentration), we should observe more GTP incorporation at MT ends. To test this prediction, we prepared shorter and longer seeds by polymerizing GMPCPP-tubulin at two different concentrations. The experimental scheme is illustrated in **Fig. 5A** (and in **Fig. S9A**). Higher tubulin concentration favored more numerous spontaneous nucleation and therefore produced shorter seeds (R1, Fig. 5A). Conversely, lower concentration lead to longer MT seeds (R2, Fig. 5A). Alternatively, if we took longer seeds and then sheered them into shorter and more numerous ones with a 25-gauge syringe needle, we would produce seeds of different lengths depending on the number of strokes applied to the seed sample (R2’5x & R2’10x in Fig. 5A). Using these different seeds, we set up polymerization reactions in the presence of the same concentration of free tubulin-GTP dimers at 5 μM in the presence of a radioactive γ^32^P-GTP tracer. While all reactions proceeded efficiently (as evidenced by almost complete polymerization shown in sedimentation assay, **Fig. 5E-F** & **Fig. S9B-C**), we consistently observed more γ^32^P-GTP incorporations into MT polymers (pellet) in reactions containing more numerous seeds (i.e. more MT ends) while overall GTP hydrolysis did not differ as much among all reactions (**Fig. 5C-D** & **Fig. S9D-E**). This result indicated that overall polymerization did not vary much among reactions since the same concentration of free GTP-tubulin dimers were used in all reactions (**Fig. 5C** & **S9C**). On the other hand, the number of polymerizing ends extending from the seeds varies greatly depending on what type of seeds were used in the reactions, and as a result we observed more GTP incorporation into the polymers in reactions with more numerous polymerizing ends (R1 vs R1’, R2 and R2’ in **Fig. 5B-F** & **Fig. S9D**).

**Fig. 5.**
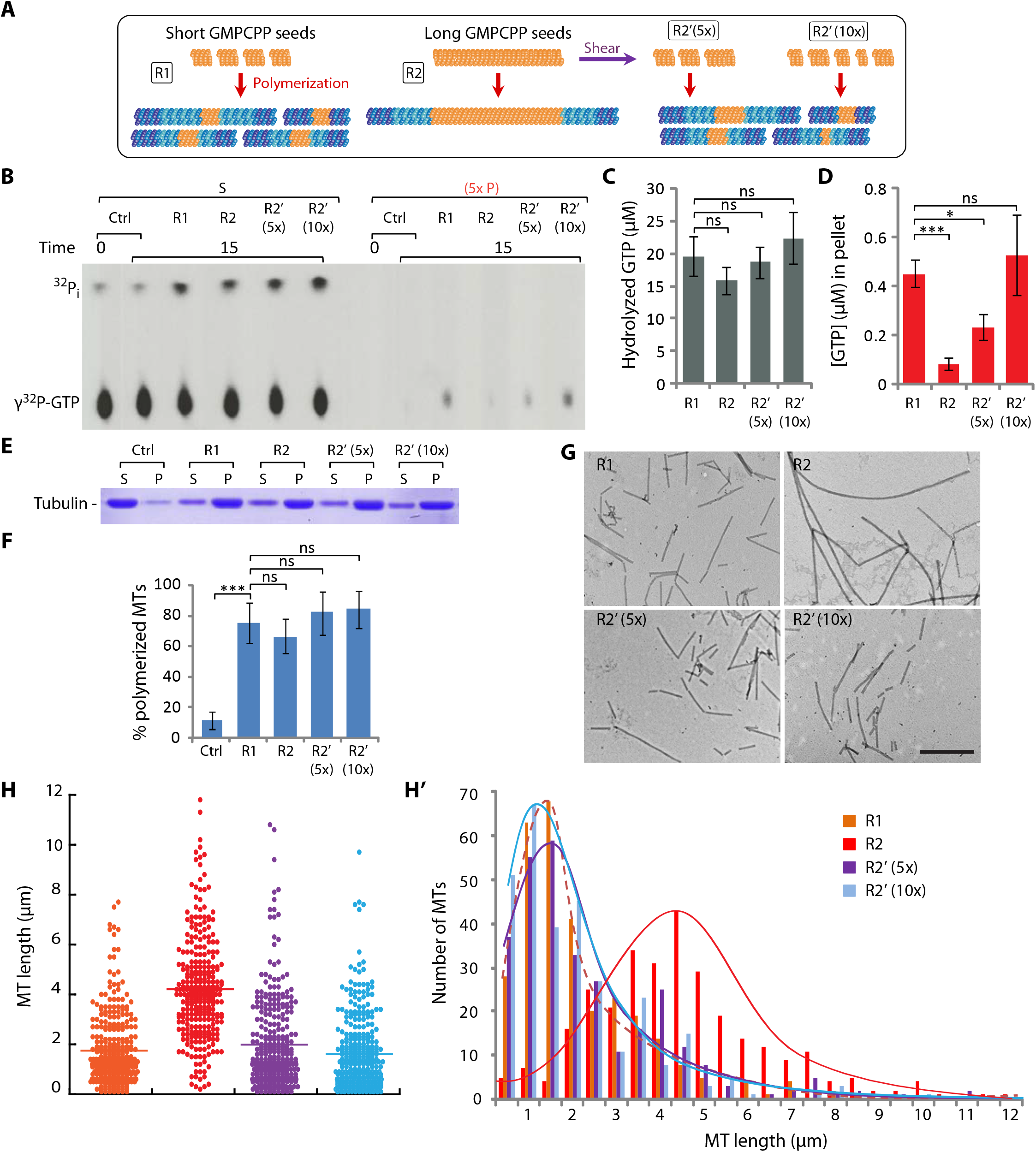
The level of GTP-bound tubulin dimers incorporation into MT polymers is directly proportional to the number of polymerizing MT ends. **(A)** An illustration depicting MTs polymerized from different GMP-CPP seeds. Briefly, MT polymerization reactions were set up with 5 μM tubulin and 200 μM GTP alone (ctrl) or in the presence of short (R1) and long (R2) GMP-CPP seeds (at 1 μM) or shorter seeds generated by shearing of long seeds through a 25-gauge needle 5 time (R2’ 5x) or 10 times (R2’ 5x). **(B-F)** GTP hydrolysis level was assessed using radio-labeled γ^32^P-GTP as a tracer during the polymerization of MTs under the indicated reaction conditions as described in (A). Samples were processed the same way as described in Fig. 4F-J. **(B)** A representative autoradiogram is shown. **(C-D)** The corresponding quantifications of overall GTP hydrolysis, as indicated by the amount of γ^32^P_i_ in the supernatant fractions (C), and of level of tubulin-GTP incorporation into MT polymers, as marked by γ^32^P-GTP in the pellet fractions (D). **(E)** The levels of MT polymerization in the reactions shown in (B) were measured using a sedimentation-based assay. A representative Coomassie-blue stained gel is shown. **(F)** The corresponding quantification of MT polymerization reactions shown in (E). **(G)** Negative stained TEM images of MTs generated from the indicated reactions as depicted in (A) and as carried out in (B). Scale bar, 1 μm. **(H-H’)** Length distributions of MTs polymerized under the indicated reaction conditions, represented by scattered plots (H) or binned histograms (H’). The lines in H indicates the mean length, while the peaks in H’ marks the median length. Data represent averages of at least 3 independent experimental sets. Error Bars, S.D. (ns (not significant): p>0.05; *p≤0.05; **p≤0.01; ***p≤0.001, by Student’s *t*-test)

To provide a quantitative account of the polymers that forms in these reactions, we determined the length of MTs by electron microscopy (EM) (**Fig. 5G**). We prepared the samples using two methods: one was to dilute the polymerized samples into paclitaxel-containing stabilizing buffer and the other was to fix it with formaldehyde, and then observed them by negative stained EM. The length measurements we obtained were consistent with what we observed in the radioactive tubulin-GTP polymerization experiments: with shorter seeds produced shorter MTs and longer seeds produced longer MTs. From these measurements and the quantification of GTP incorporation at MT ends (**Fig. 5D,F** & **Fig. 5H**), we could extrapolate the average length of GTP-tubulin-bearing protofilaments at each MT ends (= average MT length *x* % GTP in pellet, corrected by % tubulin polymerized). Based on these calculations, we determined the average length of MT ends containing GTP-tubulin to be 69.42 ± 25.53 nm (assuming 13 protofilaments/MT). This is in agreement with the length of curved protofilaments at MT ends (40-80 nm), as recently measured by McIntosh *et al.* [22] using cryo-EM images of MT polymerized *in vitro* and in various cell types. Together, these data are consistent with the idea that GTP-β-tubulin-containing MT ends reflect those dimers that have not undergone straightening and therefore still retain their conformational curvature as that of the free dimers.

### A model for conformational change driven tubulin-GTP hydrolysis

Based on the experimental data that presented thus far, we propose the follow model for conformational change driven tubulin-GTP hydrolysis (as depicted in **Fig. 6**). Free GTP-tubulin dimers exist in a native slightly curved state. When they undergo polymerization, tubulin dimers straighten up to incorporate into protofilaments within the microtubule lattice. The straightening process triggers GTP hydrolysis (as evidenced in **Fig. 4**). The polymer ends represent the transition state of this process and may retain the GTP-bound state prior to straightening (**Fig. 4F & Fig. 5**). This transition zone may represent the so-called “GTP cap” described in the literature. As tubulin dimers, free from or at MT ends, encounter kinesin-13 molecules, the complex formation further bends the tubulin dimers into an even more curved conformation, inferred from our published structural study [49]. Our experimental data show that this bending per se does not trigger GTP hydrolysis, since no hydrolysis occurs in the presence of AMP-PNP-kinesin-13s (**Fig. 1**).

**Fig. 6.**
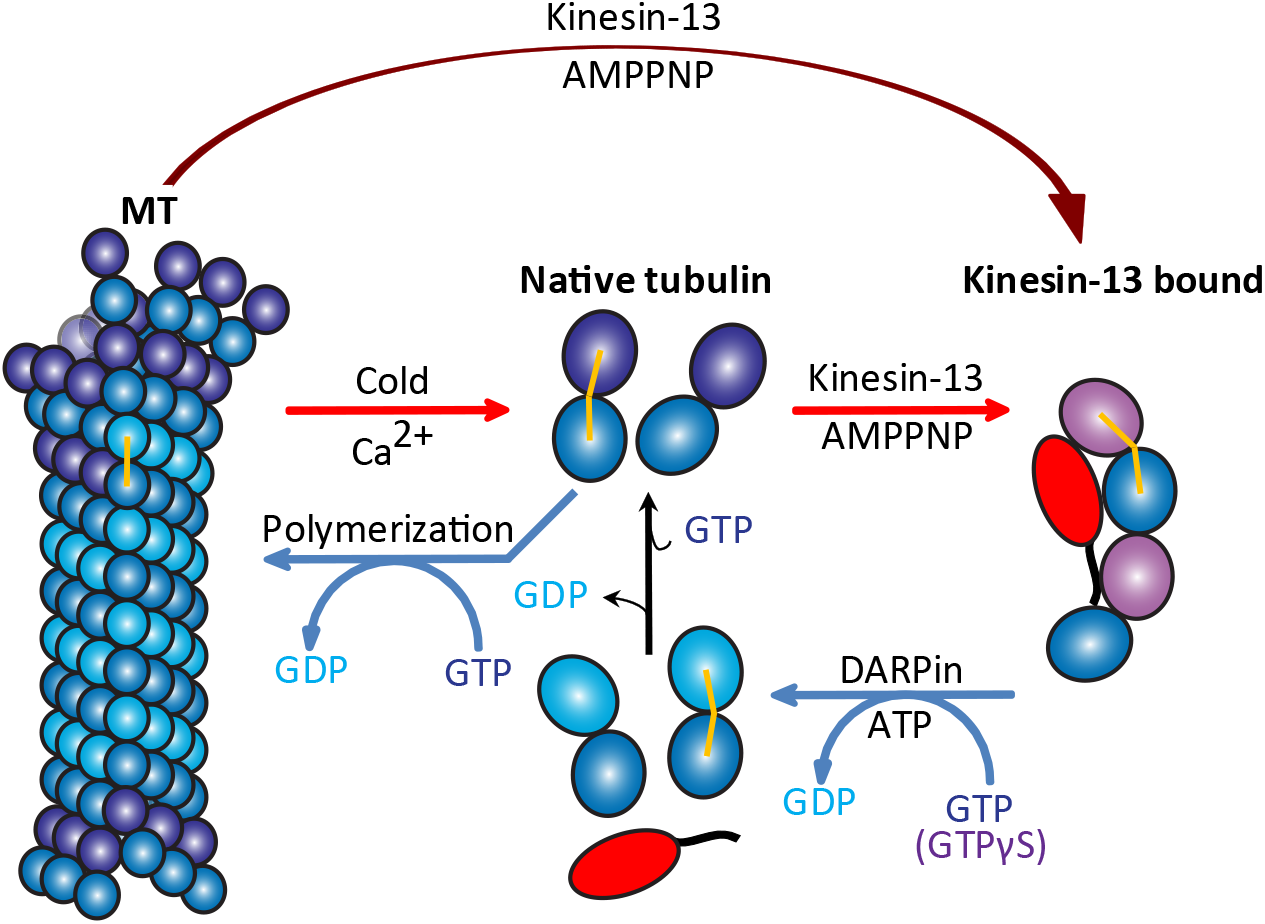
A model of conformational change-driven β-tubulin-GTP hydrolysis. An illustration depicts MTs and tubulin dimers undergoing depolymerization and polymerization. This illustration also depicts the conformational changes of tubulin dimers from their native curved (GTP or GDP-bound) to a straighter state (within MT polymer) during polymerization or to an extreme curved structure (bound to kinesin-13) during depolymerization and back to their native curvature (upon released from kinesin-13). The curvature of αβ-tubulin dimers is marked by a yellow line going through the representative tubulin dimers. Note that GTP hydrolysis only occurs when αβ-tubulin dimers undergo conformational changes from a more curved conformation to a straighter one (from right to left), but not vice versa.

However, GTP hydrolysis occurs upon the release of tubulin dimers from the complex (**Fig. 1-2**), indicating that the transition from the extremely curved state to the less curved native state triggers GTP hydrolysis, even for two less readily hydrolysable GTP analogs (GTPγS and GMP-CPP). On the other hand, kinesin-13-induced MT depolymerization differs from depolymerization mediated by Ca^2+^ or cold temperature (**Fig. 2**). The later process causes the conformational transition of tubulin from straight to the native relaxed curve, which does not trigger the hydrolysis of GTPγS. Taken together, this model presents an intriguing scenario: GTP hydrolysis only occurs when tubulin dimer transitions from curved to straighter conformations (**Fig. 6**, from right to left), but not from straight to curved (**Fig. 6**, from left to right). In addition, tubulin-GTP hydrolysis does not occur because the tubulin dimer is in a specific conformation, but it is due to the directional structural transition/change between states.

## DISCUSSION

Following our previous structural study [49], we report here that Kinesin-13 proteins induced conformational transition of associated tubulin dimers can trigger β-tubulin GTP turnover. The observation that hydrolysis of one ATP molecule can lead to hydrolysis of two GTP molecules implies that the two tubulin dimers bound to the depolymerase are both in a conformation that is more curved than the native relaxed state of free αβ-tubulin dimers and can increase their GTPase activity upon release. On the other hand, different nucleotide-bound states of β-tubulin can stimulate kinesin-13 ATPase rate to a different degree, with GTP or its mimic GTPγS-bound being the highest, followed by GDP- and Apo-state (from straighter to more curved conformations, an assessment based on the available published structural information, [54-56] (**Fig. S7A-B**). We also observed corresponding differences in binding affinity between kinesin-13 proteins and tubulin in different nucleotide states (**Fig. S7C-E**). By inference from this model, the kinesin-13 ATPase activity is highest when the depolymerase is interacting with tubulin in the shaft of a MT, as the tubulin dimers there are straightest of all and in tandem arrays. This interaction does not, however, lead to depolymerization. At MT ends, kinesin-13 proteins can interact with tubulin dimers on the exposed protofilament ends leading to the bending of tubulin dimers and spraying outward of protofilaments to facilitate its depolymerization. This is accompanied by the hydrolysis of tubulin-GTP dimers at MT ends upon their dissociation from kinesin-13 molecules, leading to the removal of GTP-tubulin-containing protofilament tips. Results form our experiments with the use of MTs formed by slowly hydrolyzable GTP analogs (GTPγS and GMPCPP, in **Fig. 2A-C**, **Fig. S4A-C** & **Fig. S5A-C**), support this assertion.

What we did not expect in the beginning was that our investigation on kinesin-13 mediated MT depolymerization would reveal something more fundamental, namely the molecular basis of β-tubulin-GTP hydrolysis. It is generally thought that GTP hydrolysis leads to a conformational change of tubulin dimers as they incorporate into MT polymer [16, 23, 58]. Our data suggest otherwise. Our experiments with paclitaxel showed that it is the change of tubulin conformation that triggers GTP hydrolysis (Fig. 4). In addition, our data on tubulin-GTP with AMP-PNP-bound kinesin-13s and DARPin also indicate that it is the change/transition of conformation, and not a particular nucleotide-bound state, that facilitates the GTP hydrolysis (Fig. 1-2). Furthermore, this change needs to be directional (from a more curved to a less curved conformation, but not vice versa) to be productive, as we have shown the lack of GTP turnover with calcium or cold induced depolymerization (from a straight to a more curved conformation). It also appears that the degree of structural change matters too. For example, GTPγS- and GMPCPP-bound tubulins, unlike GTP-tubulins do not undergo hydrolysis during polymerization. There are at least two compatible explanations for that. One is that GTPγS and GMPCPP are less readily hydrolyzed than GTP. Another is that GTPγS- and GMPCPP-bound tubulins are in a straighter conformation than GTP-tubulin, as suggested by published structural data [23, 30, 31, 59]. As a result, their structural changes upon polymerization are not as great as that of GTP-tubulin and therefore not enough to trigger hydrolysis. We favor this explanation because structural change can indeed be severe enough to induce hydrolysis of GTPγS and GMPCPP, as we have seen this occurring in KIF2A-NM induced MT depolymerization (**Fig 2A-C** & **Fig. S5A-C**). Together, our data strongly support conformational change-induced tubulin-GTP hydrolysis.

Our data from *in vitro* MT polymerization provide a quantitative measurement on the length of the elusive “GTP cap”. Our conformational-based model suggests that this length is directly linked to the curvature of the tubulin dimers. It implies that this cap should reflect the outwardly tapering curvature of protofilament ends that has been commonly observed in previous EM studies [21-23]. Based on our calculation, the length of the “GTP cap” is about 70 nm, at least for MT polymerized *in vitro*. This is consistent with a recent study by McIntosh et al. which determined the length of the curved protofilament ends to be about 40-80 nm, both *in vitro* and in multiple cell types, ranging from yeast to human [22]. The agreement between these two measurements suggest that the curved tubulin-GTP dimers are added to the MT ends during polymerization and the subsequent straightening of the newly added subunits, in part by forming lateral bond with adjacent protofilaments, activates β-tubulin-GTPase activity to accelerate GTP turnover. Since nucleotide hydrolysis is a transitional process and not instantaneous, it is possible that there are still some tubulin dimers in their unhydrolyzed GTP or GDP-Pi states near the ends of the polymerizing ends of MTs, as have been visualized indirectly by the use of fluorescently labelled EB1 [36, 60].

The model that we put forth here is incompatible with the idea of GTP islands that exist deep in the middle of MT lattice as those tubulin dimers should be already in the straight conformation and therefore should be in the GDP-bound state. However, our model is not in conflict with the idea that damage can occur in the MT middle and that the transient existence of GTP-containing dimers during the repair of damaged protofilaments in the MT lattice [61]. Beside this special scenario, tubulin-GTP dimers should be restricted to the extremities on MT polymers, at least *in vitro.* Taking this a step further, our model implies another provocative idea: the existence of the GTP cap may be merely a transition state of the tubulin polymerization process; the “GTP cap” simply marks the polymerizing ends, but itself per se does not offer active stabilizing role besides keeping the tubulin-GDP lattice from being exposed. In cells, however, MT end-binding proteins such as EB1 and its associated factors (e.g. the kinesin-13 KIF2C, KIF18B, and others) can modulate MT polymerization dynamics spatially and temporally to regulate different cellular processes throughout the cell cycle. It is conceivable that these proteins may interact with tubulin dimers at MT ends and alter their conformations, and thereby modulating the extent of the GTP caps in cells. It is also conceivable that some MAPs (e.g. MT repair MAPs or tubulin modifying enzymes) could keep sections of MT middle in GTP-bound form, creating the apparent “GTP” islands observed in cells to serve specific cellular functions [26]. All in all, the intriguing ability for αβ-tubulin dimers to self-organize into polymers of hollow tubules, the underlying molecular mechanisms that drive this process, and how associated factors regulate MT dynamics in space and time throughout the cell cycle will continue to fascinate us for years to come.

## MATERIALS AND METHODS

### Materials

Tubulin, DARPin and Kinesin-13 protein constructs were prepared as previously described [49, 62]. Most biochemicals were purchased from Bioshop Canada Inc. Nucleotides were obtained from Jena Bioscience via Cedarlane (Ontario, Canada). GDP detection kit was from ProFoldin (MA, USA) and ADP-Glo™ Kinase Assay kit from Promega Corporation. Radio-labeled nucleotides (α^32^P-ATP, γ^32^P-GTP, ^35^S-GTPγS) were purchased from Perkin Elmer.

### Microtubule Polymerization

#### DMSO polymerization

A premix was prepared with 2X BRB80, 2 mM DTT, 2 mM GTP or GTPγS and 20% DMSO. Recycled tubulin was thawed on ice and then an equal volume of the premix was added to it. Polymerization was done by incubating the reaction mixture in a circulating precision water bath at 37°C for 25-30 min.

We pelleted the ^35^S-labeled GTPγS MTs through 1 ml of warm 40% glycerol in BRB80 cushion by spinning down at 90K RPM in a Sorvall TLA100 rotor for 5 min and then resuspended the MTs in BRB80 buffer containing paclitaxel.

#### Taxol polymerization

Paclitaxel polymerization reaction mixture was prepared with 10 μM recycled tubulin in the presence of 200 μM GDP, 20 μM of paclitaxel and 1 mM DTT in 1X BRB80. Polymerization was done as described above.

#### Polymerization from GMPCPP seeds

GMPCPP seeds were prepared as follows: Recycled tubulin was thawed on ice and diluted to 30 μM in 1X BRB80 with 1 mM DTT and 0.2 mM GMPCPP. The mixture was incubated on ice for 10 min and then clarified by spinning down at 90K RPM in a Sorvall TLA100 rotor for 5 min at 2°C. The supernatant was recovered, aliquoted to 5 μl and snap-frozen in liquid nitrogen and stored at −80°C for later usage. Short GMPCPP seeds were prepared by incubating the aliquoted 30 μM GMPCPP-bound tubulin mixture at 37°C in a precision circulating water bath for 15-20 min. The mix was then diluted to 100 μl using warm BRB80 + 1 mM DTT and spun down at 90K RPM for 5 min at 25°C using TLA100 rotor. The supernatant was discarded, and the pellet was re-suspended in 100 μl of warm BRB80 with 1 mM DTT. To prepare long GMPCPP seeds, the aliquoted 30 μM GMPCPP-bound tubulin mixture was diluted to 3 μM in warm 1X BRB80 + 1 mM DTT before incubating at 37°C for 15-20 min. After preparation, the long seeds could be sheared into shorter seeds by stroking the solution through a 25-gauge needle for 5 or 10 times. To polymerize microtubules from GMPCPP seeds, the prepared seeds (short/long/sheared) were added at either 1 or 0.25 μM to a polymerization mix containing 5 μM recycled tubulin and 200 μM GTP and incubated at 37°C for 25-30 min as described above.

### Tubulin / Kinesin-13 mediated nucleotide hydrolysis assays

Kinesin-13 mediated MT depolymerization or tubulin-GTP hydrolysis assays were carried out as previously described [49, 62]. Briefly, reactions were assembled in 30 μl volume in a BRB80-based buffer (80 mM PIPES, pH 6.8, 1 mM MgCl_2_, 1 mM EGTA, 75 mM KCl, 0.25 mg/mL bovine serum albumin, 1 mM DTT, and 0.02% Tween), with the indicated concentrations of tubulin dimers, microtubules, kinesin-13 protein constructs, DARPin, and nucleotides. Reactions were carried out at room temperature for the specified lengths of time. Cold- and calcium-induced MT depolymerizations were performed in a similar manner in the absence of kinesin-13 protein constructs. After the reactions (in the case of microtubule polymerization and depolymerization assays), MT polymers remaining were separated from the free tubulin dimers by an ultracentrifugation-based sedimentation in a Sorvall TLA100 rotor at 80K RPM for 5 min at 25°C. The supernatant fractions were retrieved from the sedimentation mixture and added to ¼ volume of 4× Laemmli buffer. The polymer-containing pellets were re-suspended in an equal volume of hot 1× Laemmli buffer (prepared by diluting the 4x buffer in depolymerization reaction buffer). Equal portions of the supernatant and pellet samples were resolved on SDS-PAGE. The gel was stained with Coomassie blue dye and scanned with either Epson Perfection 4990 Photo or CanoScan 5600F digital scanners. The bands were quantified using ImageJ (NIH).

### Nucleotide hydrolysis detection methods

#### Malachite green phosphate detection assay

Malachite green-based phosphate detection assay was used to measure ATP and GTP hydrolysis rate, as previously described [62]. Briefly, reactions were assembled in 30 μl volume in the same BRB80-based buffer used for the nucleotide hydrolysis assay, with the indicated concentrations of tubulin/microtubules, kinesin-13 protein constructs, DARPin, and nucleotides. Reactions were allowed to proceed for the indicated lengths of time (10–15 min). The reactions were then quenched using equal volume (30 μl) of 90 mM perchloric acid. We then added 30 μl of the quenched mixture to 40 μl malachite green reagent in a 384-well transparent plate to develop the color. After 5-10 min of incubation at room temperature, the level of phosphate generated in each well was quantified by measuring the absorbance at 620 nm using a TECAN infinite M200 PRO plate reader. A standardized calibration curve was generated using a titration of monobasic potassium phosphate (KH2PO4) in the same reaction buffer (**Fig. S10A**).

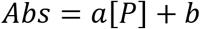

And the equation was used to calculate the concentration of phosphate generated in each well.

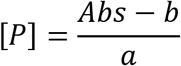

Verification of signal (or the lack thereof) from the presence of thiophosphate was done in similar manner with a titration of thiophosphate in the same reaction buffer (**Fig. S10B**). This level of background thiophosphate signal was used for background subtraction whenever applicable.

#### GDP detection assay

MicroMolar GDP assay kit (ProFoldin) was used to measure the level of GTP hydrolysis. A premix solution was prepared according to manufacturer instructions in a Tris-HCl based buffer (50 mM Tris-HCl, pH 8.0, 3 mM MgCl_2_, 0.2 mM EDTA, 0.5 mM DTT, 50 mM NaCl, 0.003% Brij-35). Equal volumes (15 μl) of reaction samples and premix solution were mixed and incubated for 45 min at room temperature. Then 30 μl of 1X Fluorescent dye was added to the mixture and after 5 min the fluorescence intensity was read at 535 nm with excitation at 485 nm using TECAN infinite M200 PRO plate reader. GDP samples of known concentrations were used to obtain a linear standard curve of the fluorescent intensity (Fc) values and the GDP concentration [GDP] (**Fig. S1A**).

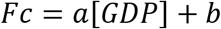

GDP concentrations were calculated using the Fc values from the unknown samples and the a and b values from the standard curve.

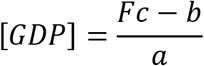

To verify the level of signal interference from the presence of ADP in the mixture, another standard curve was plotted from a separate set of assays using a titration of known ADP concentrations (**Fig. S1B**).

#### ADP Glo assay

Reactions were assembled in 25 μl total volume, as described for the Malachite Green assay. Once the reactions were complete, the ADP-Glo™ Assay was performed in two steps. The reaction mixture was then added to 25 μl of ADP-Glo™ Reagent and incubated at room temperature for 40 minutes. Then the mixture was added to 50 μl of Kinase Detection reagent (to convert ADP to ATP and introduce luciferase and luciferin to detect ATP) and incubated at room temperature for 40-60 minutes (based on the concentration of ATP). The mixture was added to a 96-well white solid bottom plate in duplicates and luminescence was measured using TECAN infinite M200 PRO plate reader. Four ATP-to-ADP conversion standard curves were prepared at different ATP+ADP concentrations (1, 10, 100 or 1000 μM), in 25μl of 1X reaction buffer. ADP-Glo™ Kinase Assays were performed at room temperature as described for the samples and four standard curves were plotted from the luminescence values of the different concentrations of ATP/ADP (**Fig. S11**).

#### Radioactivity assay

Nucleotide hydrolysis or MT polymerization assays were performed in the same buffers as described, except that the reaction mixtures were supplemented with α^32^P-ATP, γ^32^P-GTP, ^35^S-GTPγS tracers for the detection of hydrolysis. After the specified incubation time, reactions were quenched by the addition of equal volume of 1N formic acid. Small fractions of the samples were removed and spotted on a poly(ethyleneimine)-cellulose plate (EMD Millipore – TLC PEI Cellulose F – 1.05579.0001) and let air-dried for 5-10 min. Samples on the plate were resolved by thin-layer chromatography (TLC) in a glass chamber using 0.375 M KH_2_PO4 (pH 3.5) as a resolving buffer. Afterwards, the TLC plate was air dried for at least 1 hour before exposure to an X-ray film or a phosphor-imager screen. The radioactive spots on the developed x-ray films or on the scanned phospho-imager screens were quantified using ImageJ software or Typhoon FLA 9500 laser scanner, respectively. The level of hydrolysis was calculated using the ratio of the hydrolyzed spot and the initial concentration of the nucleotides (ATP_initial_ or GTP_initial_ in the solution).

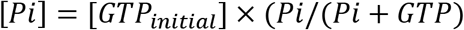

or

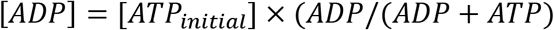

### Tubulin-Kinesin-13 Binding Assay

Tubulin-kinesin-13 binding assay was carried out via immunoprecipitation using Ni-charged MagBeads (GenScript, NJ), to pull down Histidine-tagged kinesin-13 proteins. Briefly, magnetic beads were washed with BRB80-based binding buffer (80 mM PIPES, pH 6.8, 1 mM MgCl_2_, 1 mM EGTA, 75 mM KCl, 0.25 mg/mL bovine serum albumin, 1 mM DTT, and 0.02% Tween). The reactions were started by adding kinesin-13 proteins at the indicated concentrations to 2 μl of magnetic beads in BRB80-based binding buffer. The indicated concentrations of tubulin dimers and nucleotides were then added to the reaction mixtures that were assembled in a final volume of 30 μl. Reactions were carried out at room temperature for 5 minutes. The beads were then pulled down using a magnetic stand. The supernatant fractions were retrieved from the sedimentation mixture and added to ¼ volume of 4× Laemmli buffer. The beads were re-suspended in an equal volume of hot 1× Laemmli buffer (prepared by diluting the 4x buffer in the binding reaction buffer). Equal portions of the supernatant and pellet samples were resolved on SDS-PAGE. The gel was stained with Coomassie blue dye and scanned with either Epson Perfection 4990 Photo or CanoScan 5600F digital scanners. The bands were quantified using ImageJ (NIH).

### Negative staining of MTs by Transmission Electron Microscopy (TEM)

Before sample application, the grids (Formvar Carbon Support Film on Square Grids – FCF200-Ni; Electron Microscopy Sciences) were glow discharged via a Pelco easy glow (Pelco, Fresno, USA) for 30 seconds. MT samples were prepared as described above, just right before grid preparation and were diluted to 0.5 μM. 10 μl of the diluted sample was placed on a piece of parafilm and grid was placed on the top (with carbon side touching the sample) for ~1 min. Excess liquid was then removed by gently tilting the grid sideway on a Whatman blotting paper. The grid was then rinsed with a droplet of dH_2_O and blotted again with a fresh Whatman paper. The rinse procedure was repeated two more times and the samples on the grid were then stained with 10 μl of 0.5% filtered EM-grade Uranyl Acetate (UA). After removing the excess UA by blotting, the grid was air dried for at least 1 hour. Grids were examined at the EM facility at the Université de Montréal in a FEI Tecnai 12 (Eindhoven, The Netherlands) transmission electron microscope operating at 80 kV. For each experimental condition, the lengths of 300 microtubules were quantified using ImageJ (NIH).

## Acknowledgement

We thank the LSCTB platform at the Université de Montréal for the access to the Electron Microscopy facility. We thank members of the Kwok lab for their inputs and suggestions, and Drs. Richard J McIntosh and Nikita Gudimchuk for their critical reviews on the manuscript. We acknowledge the funding support from the Canadian Institutes of Health Research (to B.H.K: PJT 148982 & 152920). B.H.K. is a recipient of the Fonds de recherche du Québec – Santé (FRQS) Chercheure-boursière Junior 1 and Junior 2 Awards and the Canadian Institutes of Health Research (CIHR) New Investigator Award. M.P. is supported by a FRQS doctoral fellowship and by doctoral and financial programs at the Université de Montréal.

## Author information

Correspondence and requests for materials should be addressed to.

## Author contributions

B.H.K. conceived the project. M.P. and B.H.K. designed and performed experiments. M.P. and B.H.K analyzed and interpreted data. B.H.K. wrote the first draft. M.P. and B.H.K. prepared and completed the manuscript.

## Abbreviations

NM: Neck+Motor
MD: Motor domain
MT: Microtubule

