## Supplemental Figures for "Evidence for conformational change-induced hydrolysis of β-tubulin-GTP"

### Figure S1

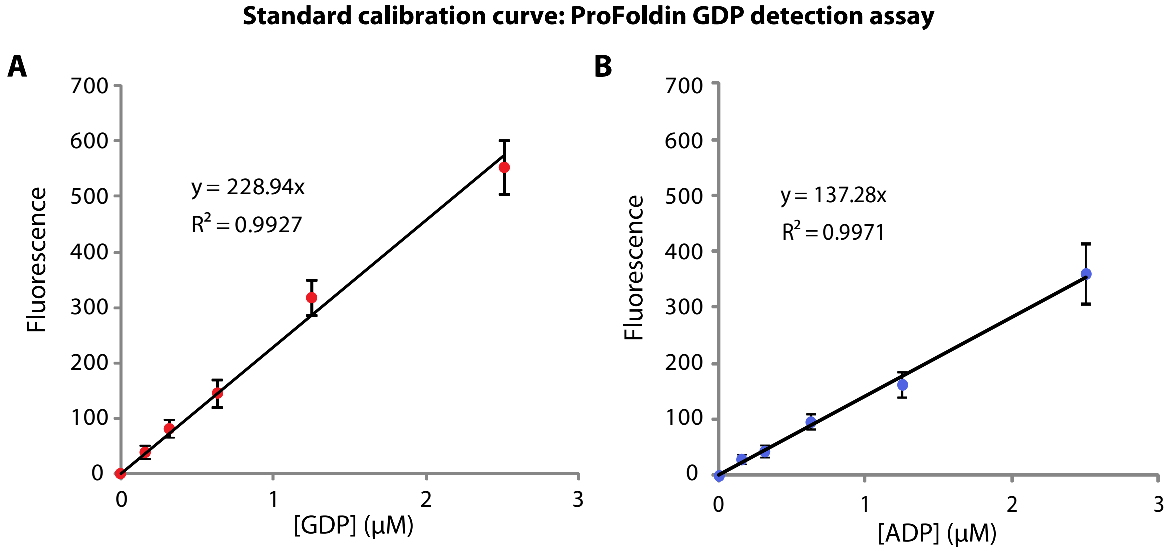


Figure S1: **Standard curves of ProFoldin MicroMolar GDP assay.** (A) Standard curve of ProFoldin MicroMolar GDP assay, using different concentrations of GDP based on fluorescence measurements with emission at 535 nm and excitation at 485 nm. (B) Similar calibration curve was generated for the assay, using a series of concentrations of ADP. Data represent averages of 3 independent experimental sets. Error Bars, S.D.

### Figure S2

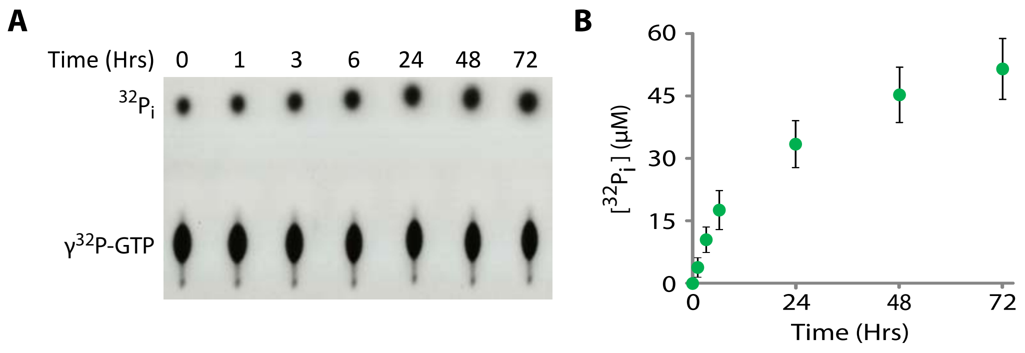


Figure S2: **Basal rate of tubulin-GTP hydrolysis.** (A) A time course experiment in which basal tubulin-GTP hydrolysis was measured using a $\gamma^{32}$P-GTP radio-labeled GTPase assay in the presence of 4 μM tubulin dimers and 0.2 mM of GTP. A representative autoradiogram is shown. (B) Quantification of $^{32}P_i$ of each time point in the time course experiment shown in (A). Data represent the average of at least three independent experimental runs. Error bars, S.D.

### Figure S3

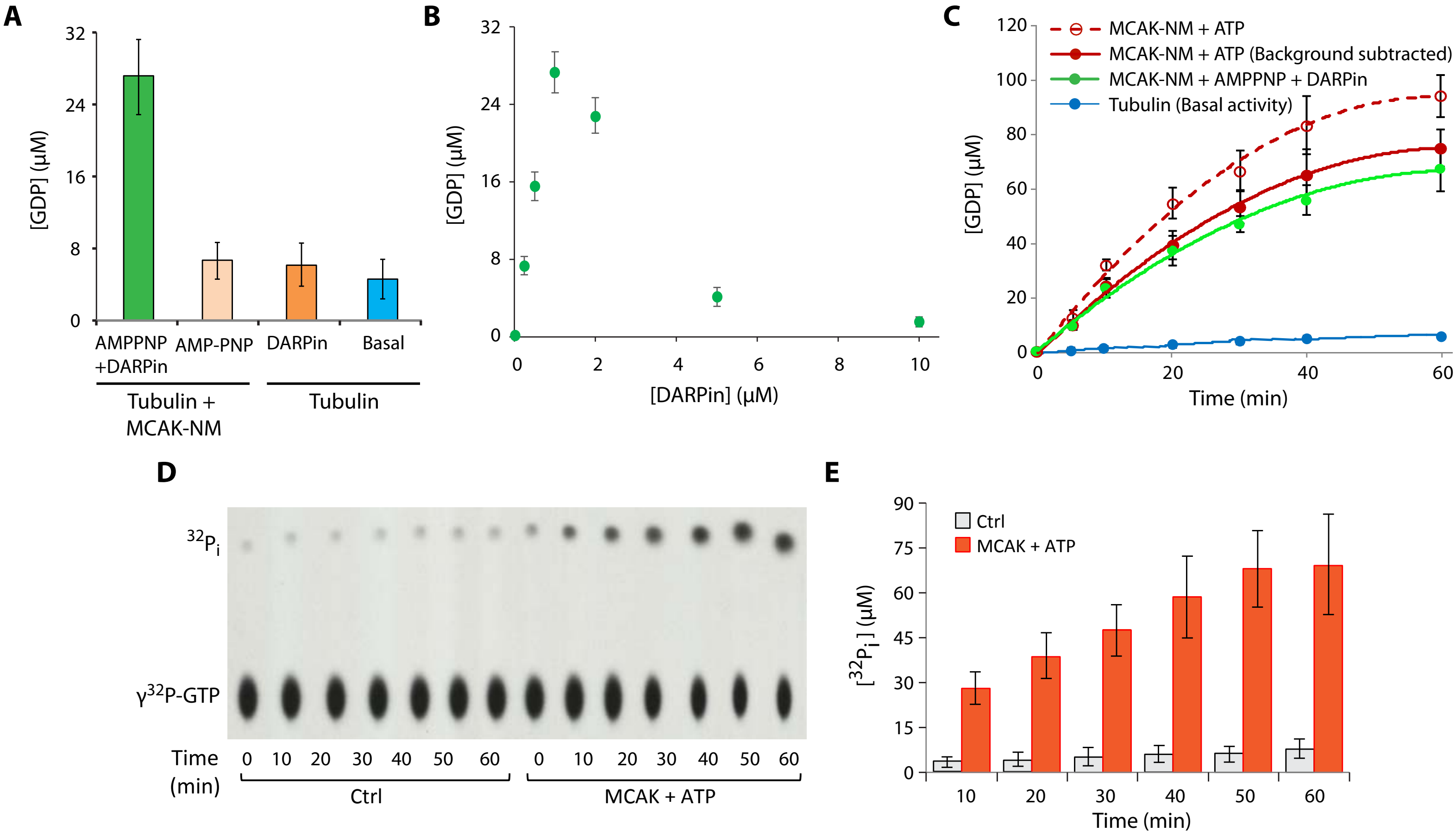


Figure S3: **The binding and unbinding of the Kinesin-13 protein MCAK to and from tubulin dimers induce β-tubulin-GTP hydrolysis.** (A) Tubulin-GTP hydrolysis measured by GDP production (ProFoldin GDP detection assay) when tubulin dimers (4 μM) and GTP (0.2 mM) were incubated alone, with DARPin or in the presence of MCAK-NM (200 nM) with AMP-PNP or with AMP-PNP and DARPin for 15 minutes at room temperature. (B) To determine the effect of DARPin, tubulin-GTP hydrolysis measured by GDP production when tubulin-GTP dimers were incubated in the presence of MCAK-NM and AMP-PNP with different concentrations of DARPin, under the same condition as in (A). (C) Time-course experiments comparing the tubulin-GTP turnovers between tubulin dimers that were incubated in the presence of MCAK-NM with AMP-PNP and DARPin and those with MCAK-NM with ATP under the same condition as in (A). For (A-C), the level of GDP was measured using a GDP detection assay (ProFoldin). (D) A time course experiment in which basal and MCAK-NM-induced β-tubulin-GTP hydrolysis (in the presence of ATP) using a $\gamma^{32}$P-GTP radio-labeled GTPase assay under the same experimental setting as described as in (C). $\gamma^{32}$P-GTP and $^{32}P_i$ were resolved by thin layer chromatography and exposed to a film. A representative autoradiogram is shown. (E) Quantification of $^{32}P_i$ of each time point in the time course experiment shown in (D). All the data shown represent averages of at least three independent experimental runs. Error bars, S.D. (ns (not significant): $p>0.05$; *$p\leq0.05$; **$p\leq0.01$; ***$p\leq0.001$, by Student's *t*-test)

### Figure S4

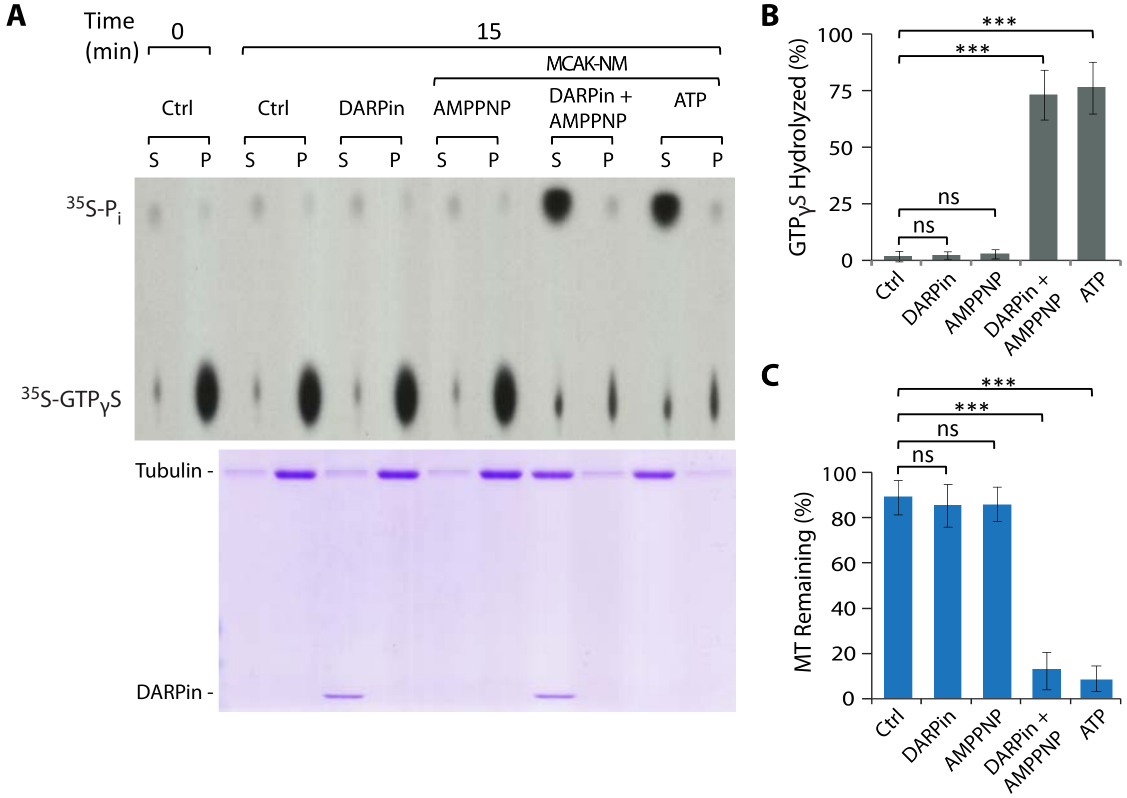


Figure S4: **MCAK-NM mediated MT depolymerization triggers tubulin-GTP hydrolysis.** (A) MT depolymerization assay was set up using $^{35}$S-GTPγS-labeled MTs alone (Ctrl) or in the presence of DARPin alone, with MCAK-NM and AMP-PNP, with MCAK-NM, AMP-PNP and DARPin, or with MCAK-NM and ATP. Reactions were carried out at room temperature for 15 minutes. Samples containing $^{35}$S-GTPγS and $^{35}$S-$P_i$ were resolved by thin layer chromatography and radioactivity was detected by exposure to a film. A representative autoradiogram was shown on the top panel. The level of MT polymers was monitored at the 15-minute time point using an ultracentrifugation-based sedimentation-based assay. Samples from the supernatant (S) and pellet (P) fractions were resolved by SDS-PAGE and the gel stained by Coomassie blue. A representative gel is shown on the bottom panel. MTs were used at 2 μM, MCAK-NM at 50 nM and DARPin at 1μM. (B,C) Quantification of data from experiments shown in (A).

### Figure S5

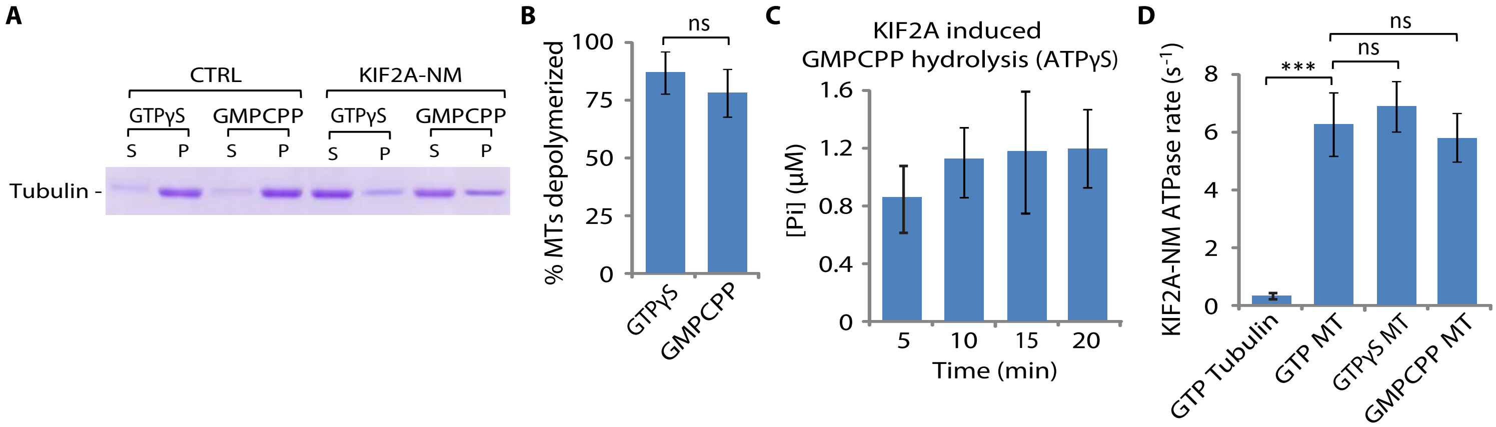


Figure S5: **Kinesin-13-mediated MT depolymerization triggers hydrolysis of β-tubulin-GMPCPP.** (A) GTPγS and GMPCPP MTs were prepared using 10 µM tubulin and 200 µM of GTPγS or GMPCPP, respectively. Microtubule depolymerization reactions were set up using 2 µM GMPCPP MTs, without (Ctrl) or with 50 nM KIF2A-NM in the presence of 200 µM ATP. The level of depolymerization was monitored using a sedimentation-based assay. Reactions were carried out at room temperature for 15 minutes. A representative Coomassie-blue stained gel from 3 independent runs is shown. (B) The corresponding quantification of MT depolymerization reactions described in (A). (C) The occurrence of GMPCPP hydrolysis during KIF2A-NM-mediated microtubule depolymerization was detected by the presence inorganic phosphate ($P_i$) using Malachite Green-based phosphate detection assay. Depolymerization reaction was set up in a time course experiment using2 µM of pre-clarified GMPCPP MTs, 50 nM KIF2A-NM and 200 µM ATPγS. Pre-clarified GMPCPP MTs were prepared by pelleting in an ultracentrifuge-based sedimentation and then re-suspending in a BRB-80-based buffer right before the experiment. The use of ATPγS, instead of ATP, here was to ensure that the signal detected was from GMPCPP hydrolysis since thiophosphate generates minimal background signal with malachite green reagent (see calibration curve in **Fig. S10B**). Reactions were carried out at room temperature for the indicated lengths of time. (D) Tubulin or MT-stimulated ATPase rates of KIF2A-NM were quantified using ADP Glo Kinase assay. Reactions were assembled, as described in the method section, with 50 nM KIF2A-NM and 200 µM ATP in the presence of 2 µM upolymerized tubulin, paclitaxel-stabilized GDP MTs, GTPγS MTs or GMPCPP MTs. Reactions were carried out at room temperature for 15 minutes. Data represent the average of at least 3 independent experimental sets. Error Bars, S.D. (ns (not significant): $p>0.05$; *$p\leq0.05$; **$p\leq0.01$; ***$p\leq0.001$, by Student's *t*-test)

### Figure S6

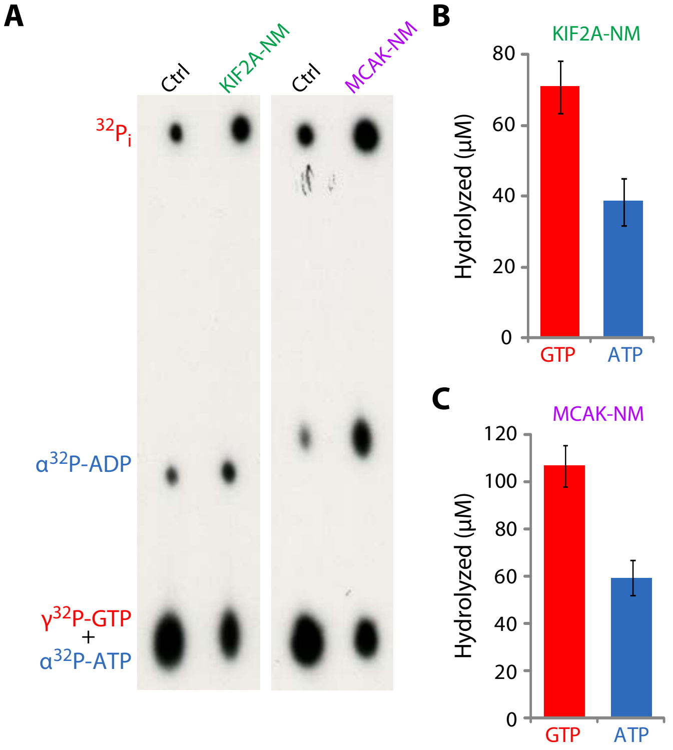


Figure S6: **Stoichiometric relationship between MCAK-NM ATPase activity and the induced tubulin-GTP turnover.** (A) ATP hydrolysis of MCAK-NM and tubulin-GTP turnover were monitored using radio-labeled $\alpha^{32}$P-ATP and $\gamma^{32}$P-GTP as tracers in an enzymatic assay, in a similar experiment as shown in Fig. 3. However, in this experiment, both radio-labeled tracers were added to the same reaction mixtures. A representative autoradiogram of a TLC plate is shown. Tubulin dimers were used at 4 µM, KIF2A-NM and MCAK-NM at 500 nM, ATP and GTP at 200 µM. Reactions were carried out at room temperature for 10 minutes. (B-C) Quantification data for the experiments shown in (A). The levels of ATP and GTP hydrolysis were quantified based on the percentage of $\alpha^{32}$P-ADP (from $\alpha^{32}$P-ATP) and $^{32}P_i$ (from $\gamma^{32}$P-GTP) of the total amount of $\alpha^{32}$P-ATP and $\gamma^{32}$P-GTP used in the corresponding reaction. Data represent the average of at least 3 independent experimental sets. Error Bars, S.D.

### Figure S7

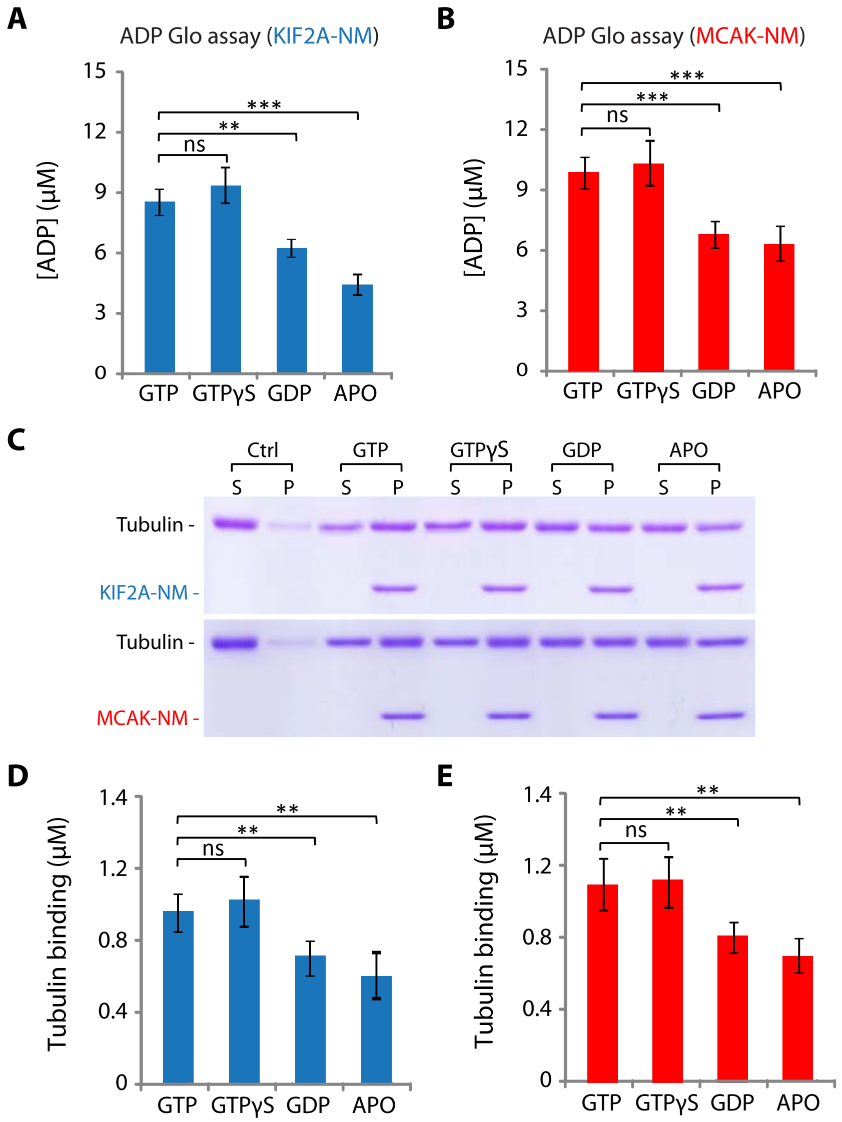


Figure S7: **Tubulin dimers in different nucleotide states differentially stimulate ATPase rates of kinesin-13 proteins and exhibit differential binding affinity to these kinesins.** (A,B) ATP hydrolysis of KIF2A-NM (A) and MCAK-NM (B) in the presence of tubulin dimers with different nucleotides as indicated using ADP Glo$^{TM}$ reagent which monitors the ADP level in the reactions. (C) Binding affinity Kinesin-13 proteins to different nucleotide-bound tubulin dimers was measured by His-tagged affinity pull down assay with Ni-coated magnetic beads. The level of tubulin dimers associated with His-tagged KIF2A-NM or MCAK-NM, was assessed by SDS-PAGE and Coomassie-blue staining. Representative gels are shown. (D-E) Corresponding quantifications of the binding data in (C) for KIF2A-NM (D) or MCAK-NM (E) are shown. Data represent averages of 3 independent experimental sets. Error bars, S.D. (ns (not significant): p>0.05; *p≤0.05; **p≤0.01; ***p≤0.001, by Student's *t*-test)

### Figure S8

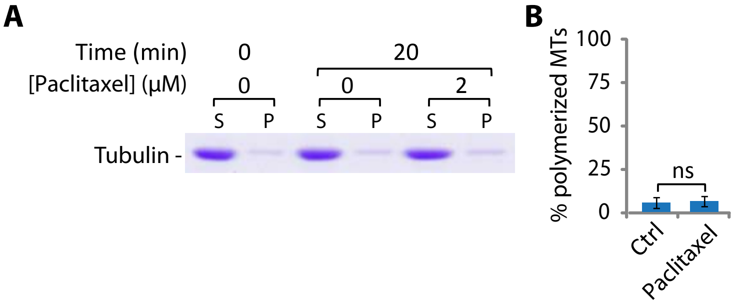


Figure S8: **Absence of MT polymerization in the presence of low tubulin and paclitaxel concentration at 4°C.** (A) The levels of MT polymerization in the reactions shown in Fig. 4A were assessed by ultracentrifugation-based sedimentation assay. A representative Coomassie-blue stained gel with negligible level of pelleted polymer is shown. (B) The corresponding quantification of MT polymerization reactions shown in (A). Data represent averages of at least 3 independent experimental sets. Error Bars, S.D.

### Figure S9

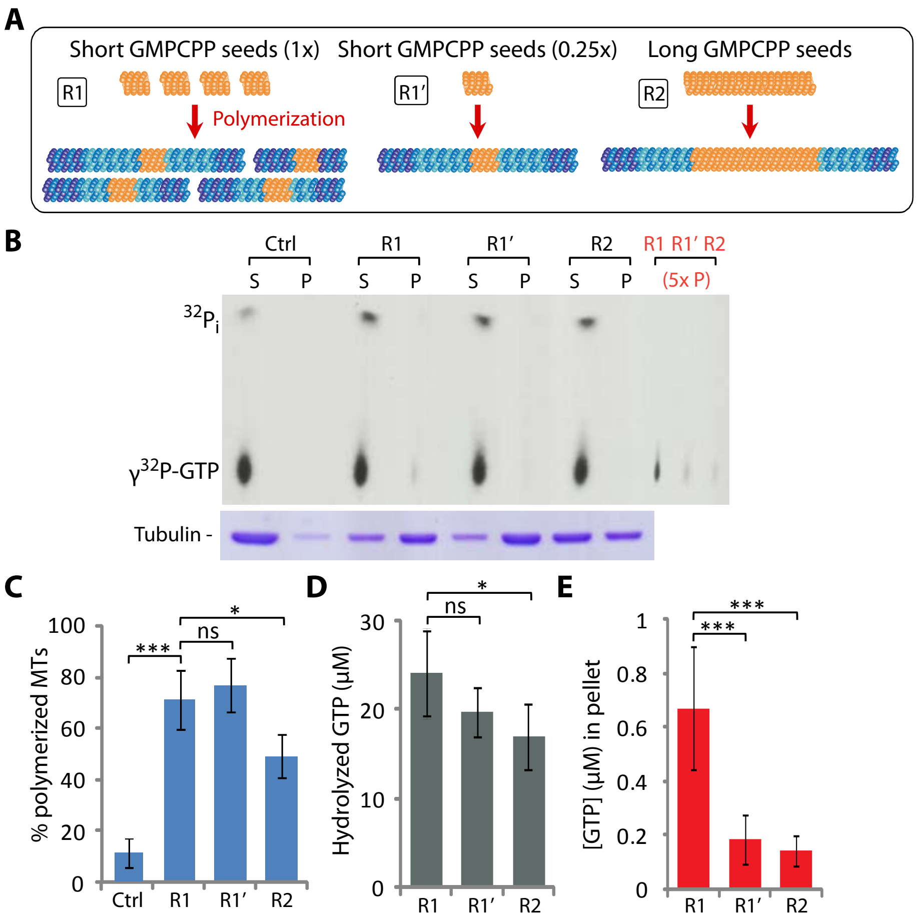


Figure S9: **The level of GTP-bound tubulin dimers incorporation into MT polymers is directly proportional to the number of polymerizing MT ends.** (A) An illustration depicting MTs polymerized from different GMP-CPP seeds, as described in Fig. 5. Briefly, MT polymerization reactions were set up with 5 µM tubulin and 200 µM GTP alone (ctrl) or in the presence of 1 µM short (R1), 0.25 µM short (R1'), or 1 µM long (R2) GMP-CPP seeds. (B-E) Level of GTP hydrolysis was assessed using radio-labeled $\gamma^{32}$P-GTP as a tracer during the polymerization of MTs under the indicated reaction conditions as described in (A). Samples were processed the same way as described in Fig. 4F-J. Note that the pellet (P) fractions were loaded at 5x equivalence of the amount of the corresponding supernatant (S) fractions, in order to detect the level of γ32P-GTP more readily. (B) A representative autoradiogram is shown (top). The levels of MT polymerization were measured using a sedimentation-based assay. A representative Coomassie-blue stained gel is shown (bottom). (C) The corresponding quantification of MT polymerization reactions shown in (B). (D-E) The corresponding quantifications of overall GTP hydrolysis, as indicated by the amount of $\gamma^{32}P_i$ in the supernatant (S) fractions (D), and of level of tubulin-GTP incorporation into MT polymers, as marked by $\gamma^{32}$P-GTP in the pellet (P) fractions, corrected by the loaded amount (E). Data represent averages of at least 3 independent experimental sets. Error Bars, S.D. (ns (not significant): $p>0.05$; *$p\leq0.05$; **$p\leq0.01$; ***$p\leq0.001$, by Student's *t*-test)

### Figure S10

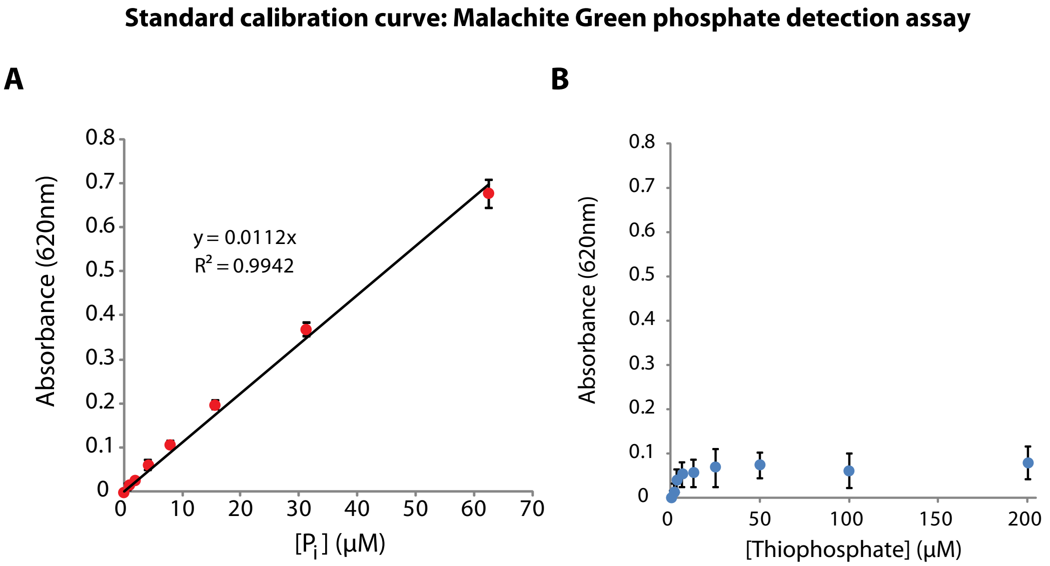


Figure S10: **Standard curves of Malachite Green-based phosphate detection assay.** (A-B) Standard calibration curves of Malachite Green-based colorimetric phosphate detection assay, generated by measuring different concentrations of inorganic phosphate (A) or thiophosphate (B) at absorbance of 620nm. Data represent averages of 3 independent experimental sets. Error Bars, S.D.

### Figure S11

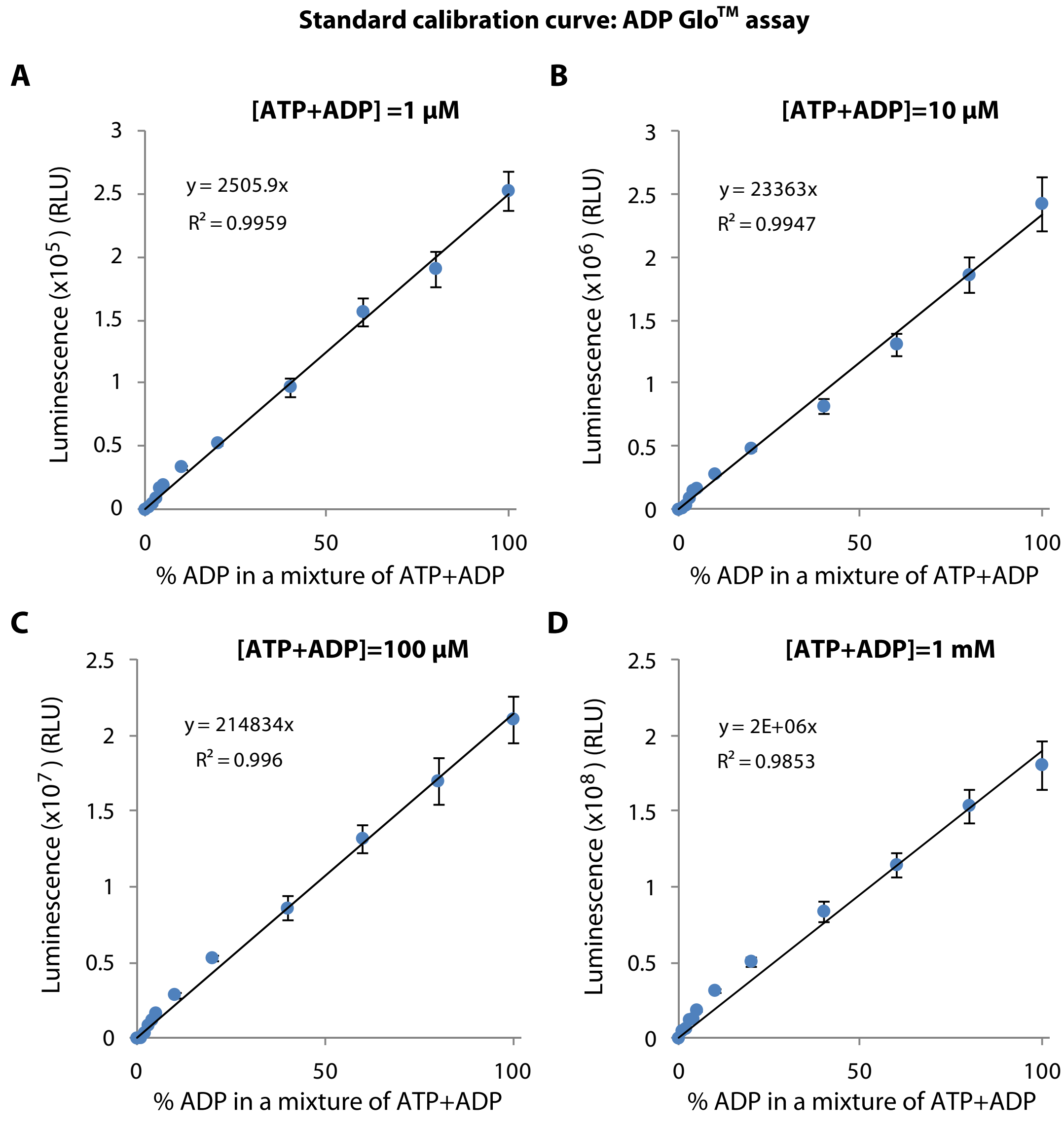


Figure S11: **Standard curves of ADP-Glo™ Kinase assay.** (A-D) ATP-to-ADP conversion curves, using ADP-Glo™ Kinase assay, at the indicated ATP+ADP concentrations, 1 µM (A), 10 µM (B), 100 µM (C) and 1 mM (D). Plots showlinear fits of luminescent signal and the amount of ADP in the reaction mixtures of ATP+ADP.
